# Plague risk in the western United States over seven decades of environmental change

**DOI:** 10.1101/2021.02.26.433096

**Authors:** Colin J. Carlson, Sarah N. Bevins, Boris V. Schmid

## Abstract

After several pandemics over the last two millennia, the wildlife reservoirs of plague (*Yersinia pestis*) now persist around the world, including in the western United States. Routine surveillance in this region has generated comprehensive records of human cases and animal seroprevalence, creating a unique opportunity to test how plague reservoirs are responding to environmental change. Here, we develop a new method to detect the signal of climate change in infectious disease distributions, and test whether plague reservoirs and spillover risk have shifted since 1950. We find that plague foci are associated with high-elevation rodent communities, and soil biochemistry may play a key role in the geography of long-term persistence. In addition, we find that human cases are concentrated only in a small subset of endemic areas, and that spillover events are driven by higher rodent species richness (the amplification hypothesis) and climatic anomalies (the trophic cascade hypothesis). Using our detection model, we find that due to the changing climate, rodent communities at high elevations have become more conducive to the establishment of plague reservoirs—with suitability increasing up to 40% in some places—and that spillover risk to humans at mid-elevations has increased as well, although more gradually. These results highlight opportunities for deeper investigation of plague ecology, the value of integrative surveillance for infectious disease geography, and the need for further research into ongoing climate change impacts.

## Introduction

The distribution and burden of infectious diseases will be entirely reshaped by global environmental change. Scientific consensus suggests that over the next century, the combined effect of climate change, land degradation and transformation, and increasing human-wildlife contact will bring about a massive increase in the spillover of pathogens that originate in wildlife (zoonotic diseases)^1,2^ and the burden of infections transmitted by arthropods (vector-borne diseases)^3,4^. While there is substantial research efforts working to project these future changes, the impacts of current environmental change on infectious disease burden in the world today is underexplored. Based on current evidence, land use change is the best-supported leading driver of zoonotic emergence^5,6^; much less is known about climate change impacts to date. This is due, in large part, to methodological limitations: the “detection and attribution” methods that are best suited to this problem require substantial data on disease prevalence or incidence over extensive periods, as well as complicated model designs (e.g., counterfactual climate scenarios without climate change)^7,8^.

Instead, many projections of climate change impacts rely on *ecological niche models* (also known as species distribution models), a set of regression and machine learning approaches that relate climate to the geographic range of a species^9,10^. Usually, these approaches are an oversimplification of reality, especially for pathogens: for example, a map of anthrax (*Bacillus anthracis*) may classify west Texas as an endemic zone, even though the system is characterized by epizootics that are sometimes years apart^11^. Ecological niche models are therefore an imperfect tool for exploring climate change impacts. These methods work well for mapping current distributions, for projecting single-time-slice distributions under future climates, and – in some recent work – for projecting continuous-time change^12^. Retrospective work to reconstruct climate change impacts is much rarer, and is usually restricted to work that builds two species distribution models for contrasting time intervals and compares them^13,14,15^. This approach is less than ideal, forcing researchers to violate the assumption that species’ geographic ranges are at equilibrium^16,17^; to aggregate data into somewhat arbitrary time periods; and to compare models trained on non-independent but non-overlapping datasets, which will generate different biological response curves simply because of model uncertainty. In this framework, it is also difficult to eliminate alternate hypotheses for why a species’ apparent distribution might change, like noise in the detection process or shifting abundance patterns.

Recently, a growing set of tools have tried to grapple with the temporal variability exhibited by the distribution of infectious diseases. Though most disease maps are treated as the long-term average of temporally-dynamic processes, *time-specific ecological niche modeling* has been proposed as an alternative that captures the dynamic nature of transmission. Almost always, though, these methods have been implemented at the finest temporal scales: monthly^18^ or seasonal^19,20^. As of yet, this approach has been mostly untested as a way of understanding disease distributions over multiple years—and ideally, of contextualizing the impacts of environmental change over decades (but see^15^).

Few systems provide a better opportunity to test this approach than plague, a globally-cosmopolitan zoonotic infection caused by the bacterium *Yersinia pestis*. The global distribution of plague has been far from stable over the past two centuries; the Third Pandemic (late 18^*th*^ to mid-20^*th*^ Century)^21,22,23,24,25^ in particular was responsible for the introduction of *Y. pestis* into many new regions that were environmentally suitable but otherwise uncolonized, particularly the Americas^26,27,28^. In some of these regions, outbreaks have faded over time, while in others, plague foci have persisted and the pathogen has become endemic, maintained by a sylvatic cycle in rodent reservoirs and flea vectors^28^. Rodent biodiversity hotspots may be particularly conducive to the formation of these reservoirs^29^, a possible case of *biodiversity amplification* effects^30,31^, where the diversity of competent hosts allows a virulent pathogen to be maintained at more stable levels. Though underexplored, emerging evidence also suggests that plague may persist in the soil, possibly by acting as an endosymbiont with amoebas^32^, from which it sporadically can reinfect burrowing rodents^33^. Soil conditions may therefore further constrain the distribution of plague reservoirs^34,35,36,37^. Like other pathogens that can persist in the soil^38,39^, provisionary evidence suggests that plague may be limited by soil salinity^40,41^, soil organic carbon, and alkalinity^42^. Though these factors may have limited influence in the short-term dynamics of plague in any one location, at continental scales, they could reasonably be expected to shape where plague foci have become established.

Both experimental and ecological analysis suggests that plague dynamics are also highly sensitive to climatic conditions. The disease’s sensitivity to bioclimatic conditions has been documented throughout its life cycle, but is particularly pronounced on the arthropod level, where temperature (and to a lesser degree humidity) influence the rate at which various flea species move through their life cycle^43^. Flea species differ in their temperature sensitivities^43^, making the local composition of flea communities an important consideration, as well as in their ability to transmit plague, either through early-phase transmission or blockage-induced transmission^44,43,45^. The bacterium appears able to rapidly evolve its ability to favor one over the other transmission mode^46^ (or maintains a standing variation in its extended phenotype within populations). Temperature also directly influences biochemical aspects of the transmission efficiency of the plague bacterium, particularly when temperatures rise above 27°C^44^, presumably by negatively influencing the stability of the biofilm that the bacterium forms in fleas. Temperature also finally influences rodent populations, including through a mechanism generally referred to as a *trophic cascade*: climatic anomalies influence primary productivity, driving changes in rodent density, which in turn change the density and biting preferences of fleas^47,48^. The combination of these environmental sensitivities, when playing out across the scale of ecosystems, can lead to widespread synchronicities in plague epizootic periods^49^.

All of these lines of evidence suggest that plague should be broadly sensitive to environmental change, and that in systems where trends in plague occurrence have been tracked, an anthropogenic signal might be detectable. The United States is the perfect system to test this approach, as data in this region are particularly abundant; human case data goes back over a century, to plague’s first introduction on the Pacific coast in early 20^*th*^ Century^50,51,26^. Moreover, the U.S. Department of Agriculture has collected plague seropositivity data from wildlife for multiple decades through the USDA National Wildlife Disease Program^52^. Combined, these national datasets include more records than many global studies of pathogen distributions^38^, making this system an ideal testing ground. Together, these data also cover nearly a century of environmental change, a temporal scope that allows time-specific ecological niche modeling to be implemented. This also allows us to revisit one of the only previous attempts at this approach, which compared models of plague risk in the western U.S. based on case data in three multiyear time slices (1965-69, 1980-84, and 1995-99), and concluded that plague risk had expanded since 1950 and would continue to do so in the future^15^.

In this study, we revisit this prediction by using two independent data streams (human cases and wildlife serology) in a machine learning model called *Bayesian additive regression trees* (BART)^53^ (see Methods for a detailed explanation). Climatic reconstructions are readily available for the duration of our study (1950–2017), allowing us to use annual climate layers (including long-term anomalies) corresponding to the year of each plague case. This pairing allows us to improve model precision relative to long-term averages, to differentiate areas of ephemeral versus persistent risk, and to identify the fingerprint of environmental change in risk trends. We also test whether the distribution of plague in this region is responsive to rodent biodiversity or soil chemistry and macronutrients, offering detailed insights into the factors that maintain plague risk. Finally, we propose a new approach that harnesses *BART with random intercepts* (riBART) to account for historical variability in detection and sampling, allowing us to confidently identify the signal of changing environmental conditions in plague prevalence over time. In doing so, we propose the first extension of ecological niche modeling that nods towards the ultimate aim of detection and attribution of anthropogenic climate change impacts on the geographic distribution of infectious diseases.

## Results

### The distribution of plague

We generated two primary models of plague over time. The first covered 9,761 animals sampled for plague (2000 to 2017), and performed well (training AUC = 0.836; Extended Data Figure 1). The second covered a total of 430 human cases of plague (1950 to 2005), and performed very well (training AUC = 0.909; Extended Data Figure 2). When both models were rerun with an overlapping “test period” of 2000 to 2005 withheld, they performed adequately, with the human model (AUC = 0.820) performing better than the wildlife model (AUC = 0.775). As both models performed well in temporal cross-validation, we used both to make annual predictions from 1950 to 2017, and split predictions into binary presence or absence risk maps for each year using the true skill statistic.

Both models found that the majority of plague risk in the western United States is, as expected, found west of the 100^*th*^ meridian (Figure 1). The human model mostly predicts risk in the “Four Corners” region (Utah, Colorado, Arizona, and New Mexico), where that risk is relatively stable across years. In contrast, the wildlife model predicts risk fairly expansively west of 100°, including at much higher latitudes than the human risk model. Risk varies much more across years in this model, but several areas are predicted to be environmentally suitable across years from Montana to west Texas. The suitable areas identified by the wildlife model in the southwest are less uniform than the human model, likely reflecting a finer-scale differentiation of risk. There are two main reasons the human model might discriminate less in this region: human cases may be reported in different locations than the site of initial spillover, and occurrence points were randomly resampled at the county level (as data have been previously de-identified).

**Figure 1:**
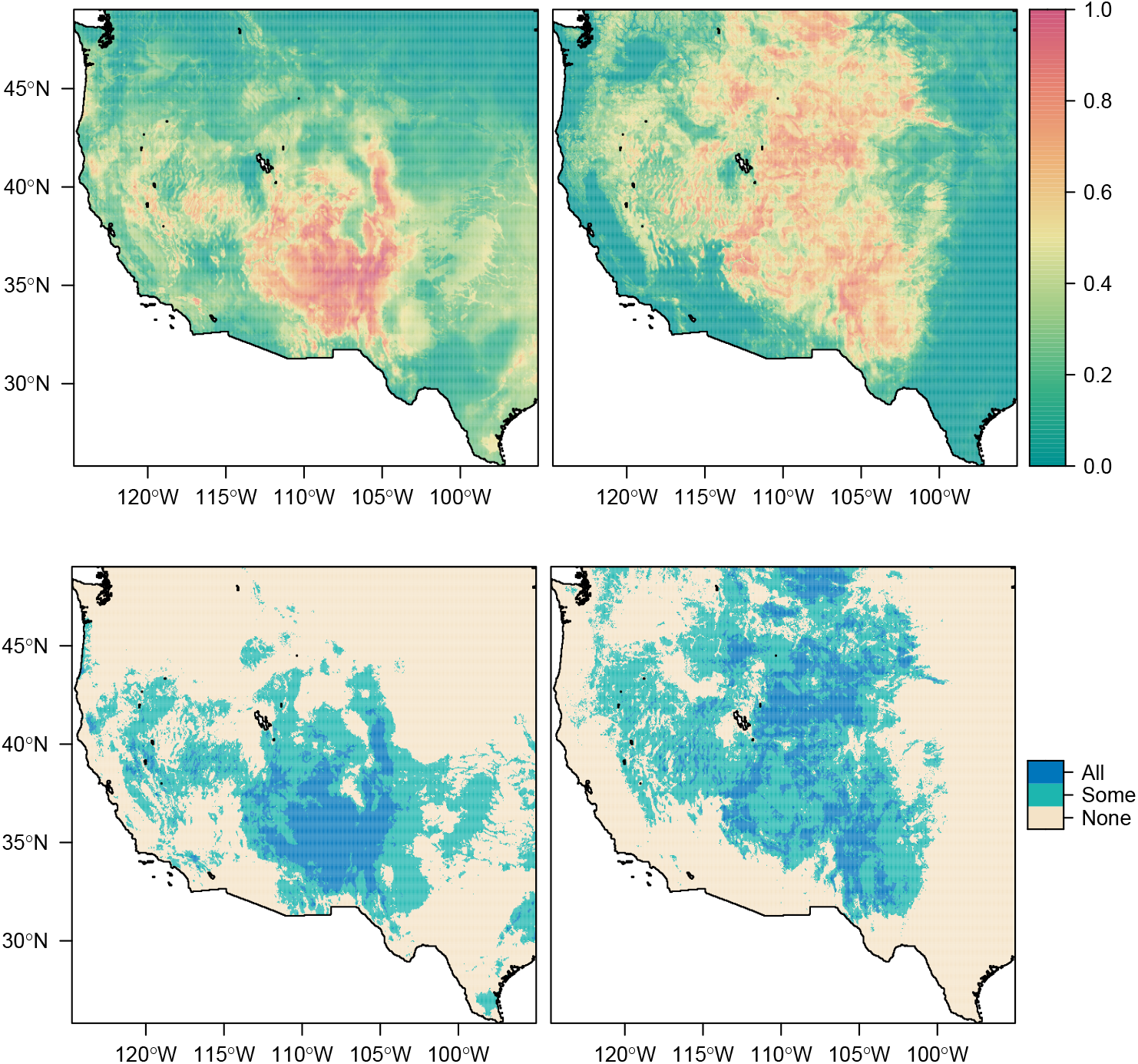
Suitability for plague across all years (1950-2017), for humans (left) and wildlife (right). Top panels give mean suitability across all years; bottom panels show areas identified as suitable in no years, at least one, or all 68 years.

For the most part, we found that risk areas identified in the human model were a subset of the much broader predictions made by the wildlife model (Figure 2), with three major exceptions. The human model identified much broader risk in southern Arizona and New Mexico, likely due to how the cases were randomized at county levels. The human model also predicted areas of risk further east, in regions like east Texas or Oklahoma where plague is not known to be endemic (and, in this regard, the wildlife model better captures the known distribution), but conditions may be broadly favorable. Finally, and most notably, the human model predicted plague risk throughout California, in places that have previously been identified as high-risk^54^. This likely reflects a deficit of data from Californian sources in our wildlife model, as state wildlife surveillance is curated independently. Together, our findings indicate the value of comprehensive surveillance, and the possibility that zoonotic reservoirs may be more expansive than areas of known spillover.

**Figure 2:**
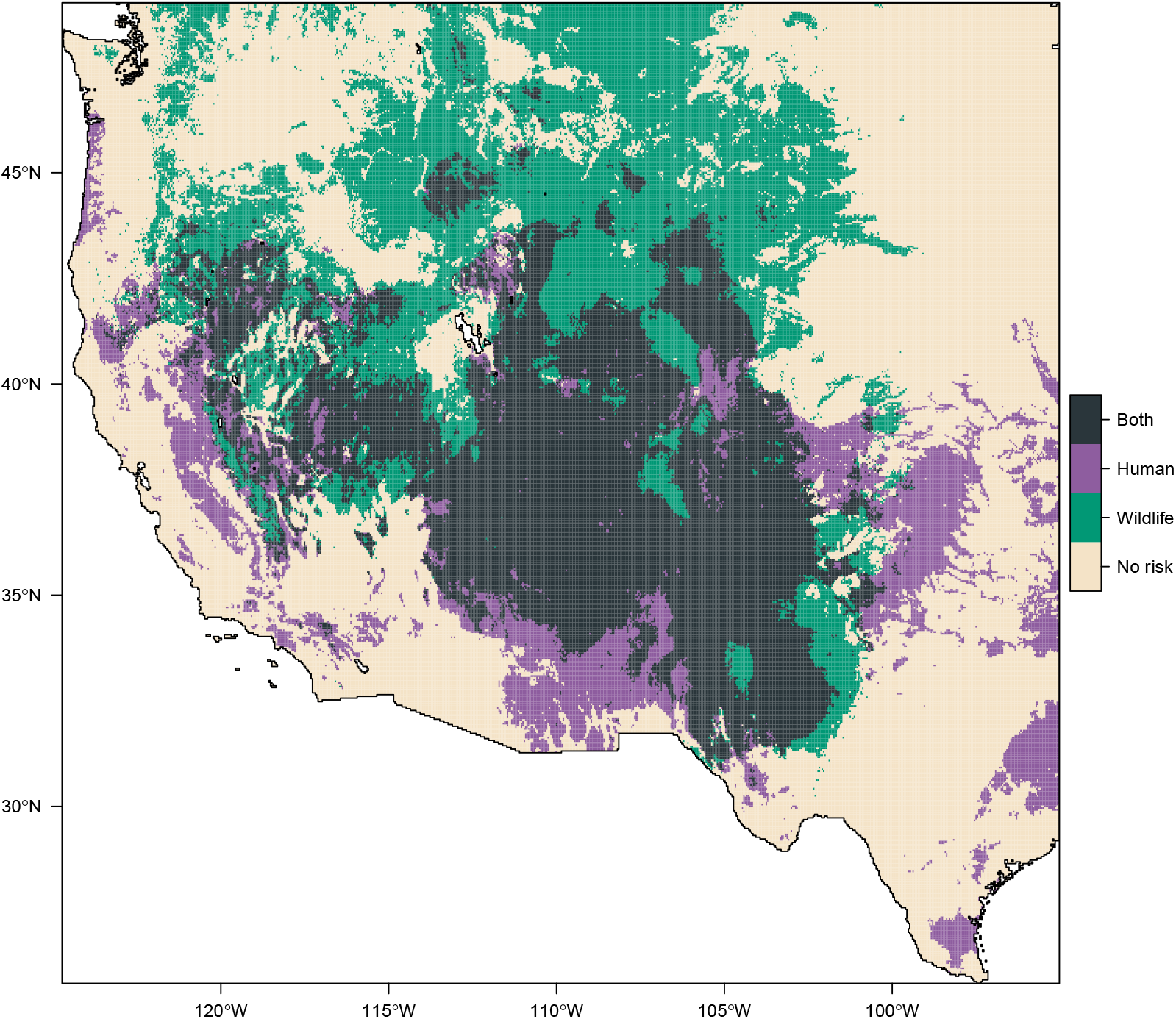
The wildlife model’s predictions largely encompass the human model’s predictions, except in southern Arizona and California (where predictions extend into other areas of suspected plague risk) and west Texas (too far east for plague reservoirs).

### Ecological insights

Our models identified a number of intersecting factors that maintain plague reservoirs and create the right conditions for spillover, many of which have been previously identified by ecological and epidemiological models (Extended Data Figures 3, 4, 5, 6, 7, 8, 9). A handful of factors are important in both animal and human spillover models, and have similar response profiles: elevation, with higher risk at higher elevations; rodent species richness, with a similar positive effect; and the sodium (Na) and calcium (Ca) content of the soil, both with a negative effect on plague risk. As these factors are shared between the models, we can tentatively conclude that these factors relate to what happens in the wildlife, and are not substantially altered by the additional spillover process that the human model incorporates. A fourth environmental factor that is significant in both models is temperature, but with different response profiles in how means, maxima and anomalies in temperature affected the risk of plague. Finally, we found strong effects of clay and iron content of the soil, which are shared between the two models but follow different profiles, as well as the sandiness of the soil (wildlife model only) and the variability in annual precipitation within the area (human model only). A list of all variables and abbreviations is given in Extended Data Table 1.

#### Elevation

Both models indicate that plague risk increases at higher elevations, particularly above 2,000 meters, compatible with previous findings in this system^54,55,56,57^. Using spatial partial dependence plots, we were able to show that the abrupt transition in plague suitability at the 100th meridian (100° W) was driven by the elevational layer in both models, and not suitably explained by any bioclimatic factors (Extended Data Figures 10, 11). Elevation has also been previously associated with plague on other continents^42,58,59,60,61,62^, and while the general trend is that there is a lower threshold elevation (and an upper limit, at the extreme altitudes in the Himalayas), that threshold differs substantially between countries. For example, Brazil’s plague reservoirs start at 500 meters above sea level, and are limited by the landscape to no more than 1,000 meters^63^, while Madagascar’s urban plague reservoir in Mahajanga is at sea level^64^, as are the plague reservoirs surrounding the Caspian Sea in Central Asia^65^. Elevation therefore seems to represent the local ecology and distributional limits of fleas and rodents, rather than a global proxy for a bioclimatic or atmospheric variable (e.g. partial CO_2_ pressure).

#### Rodent richness

Likewise, both models found a higher suitability for plague in areas with higher rodent richness, with the factor being only second to elevation in importance in the human plague-risk model. This points to the possibility that high-elevation hotspots of rodent biodiversity may help maintain enzootic plague transmission, a possible case of the biodiversity amplification effect that has also been found by similar work in China^29^. As in China, it is unclear whether the increased biodiversity itself has a positive effect on plague maintenance, or whether it merely signals an increased chance that certain key rodent species (particularly synanthropic ones) are locally present. If there are positive, general associations between rodent diversity and plague risk, this would be an exception to widespread evidence of biodiversity dilution effects for other vector-borne diseases^66^. Most theoretical models of the dilution effect rely on a skewed distribution of host competence, where higher host diversity leads to reduced transmission in the narrow subset of competent hosts^67,68,69^; plague is perhaps uniquely capable of infecting and causing disease in hundreds of host species^70^, and therefore may not exhibit these dynamics, though only a limited number of species develop a high enough viremia to infect a feeding flea. Alternately, it may be that scale underlies this pattern; theory suggests that dilution effects are strongest at small scales, while amplification effects may be normal at continental scales^30,71^. Finally, it might simply be that plague behaves differently than other vector-borne diseases because plague can also spread through pneumonic transmission and prey consumption, which produce different dynamics.

#### Climate

Temperature was a universally-important predictor across all models, while surprisingly, precipitation only minimally influenced predictions. In the wildlife model, we found a negative relationship between mean temperature and plague risk–an unusual response curve for a vector-borne disease^72^. Plague across the globe occurs in a wide variety of ecosystems, from the tropics in Africa, to semi-arid deserts in Kazakhstan and the high mountains in Central Asia, and appears able to persist across a large temperature range. Its apparent association with colder locations in the United States may therefore not be directly related to some temperature threshold, but possibly to the ecological niche of key maintenance hosts or key vector species. For human plague risk, we see a sharp increase as the mean and maximum annual temperature falls between 5° and 14°, respectively, matching previous findings from empirical work on North American fleas^73,74^ and global models of the Third Pandemic spread^75^. This may reflect the underlying thermal ecology of the pathogen^72^, or potentially some combination of rodent habitat and human behavior (e.g., these thresholds might delineate the general type of wilderness areas in the Rocky Mountains where people live alongside plague reservoirs).

In addition to the effect of long-term climatic averages, the temporal structure of the model allowed us to detect a strong effect of interannual variability. In the wildlife model, we observed an increase in plague prevalence during anomalously warm years, a result that has been previously reported for semi-arid desert ecosystems^76^, as well as for human cases in the United States^51^. Warmer years are likely to increase rodent density, both directly through mild winters^77^ and indirectly through higher primary productivity; flea populations in turn tend to follow rodent density, with some degree of lag^77,78,79^. In contrast, in the human model, spillover was most likely in anomalously wet, cold years. This matches previous findings in other systems^78,79,80^, which have been attributed to another kind of tropic cascade: when seasonal fluctuations become unfavorable to rodent populations after a recent high, and these rodent populations contract, fleas aggregate on the remaining rodents, both facilitating the dissemination of plague between rodents, and making fleas more eager to seek secondary hosts to feed on, thus leading to increased spillover risk.

#### Soil

Finally, we found that both models provided evidence that the long-term persistence of plague foci is related to properties of the soil. Our modeling suggests that *in vivo*, soils with higher proportions of sand and intermediate proportions of clay (~20-30%) (Extended Data Figure 8), low sodium and calcium contents, and mid-to-low concentrations of iron seem to be most conducive to plague. Although not included in either model after variable set reduction, we also found that soil pH may limit persistence, with more alkaline soils favored in the wildlife model. The observed response curves are somewhat unusual, given that both human and wildlife cases peaked in the raw data around a soil pH of 8.2-8.4 (Extended Data Figure 12); it may be that this reflects colinearities with other soil traits, or simply a smaller effect of pH compared to other soil characteristics.

The role of a soil compartment in the maintenance of plague reservoirs has been under consideration for more than a century, and various aspects of a soil-cycle of plague have been independently confirmed in laboratory settings. These include survival in the soil for months to years in a laboratory setting, either in association with amoebas (*Acanthamoeba castellanii* and *Dictyostelium discoideum*) or independently^34,32,36,81^; the existence of *Y. pestis* in soil in wildlife plague foci^40^; the sporadic return from soil into a rodent population^33^; and geographic correlations between plague foci and various soil properties^42,59,40^. Mechanisms through which these factors affect plague foci may be directly related to the bacterium, or through soil factors that influence the vector or the host. Fleas living in burrows, for example, are negatively affected in all aspects of their lifecycle (fecundity, development, survival, and activity) in environments with a 100-fold higher level of fractional CO_2_ than the atmospheric fraction^82^. This link to ventilation can explain the role of soil characteristics: a poorly ventilated burrow (impermeable, non-sandy soil) may reach up to 65-fold higher level of fractional CO_2_, whereas a unventilated permeable (sandy) soil would only reach up to 25-fold higher levels, and a well-ventilated permeable soil would only reach up to 7-fold higher^83^. Likewise, soil mineral content could have downstream effects on the homeostasis of virulence-related minerals in the body of the host. As a parallel, one study in Brazil found that cattle grazing on iron-rich soil would lead to highly lethal infections of normally non-pathogenic *Yersinia pseudotuberculosis*, presumably by perturbing the ability of the host to bind iron away from being utilized by the bacterium during an infection^84^. Both iron^85,86,87^ and calcium^88,89^ are virulence factors of *Y. pestis*, and we tentatively hypothesize that plague foci persist better in regions where *Y. pestis* is not facilitated by the environment to be highly virulent.

### Detecting environmental impacts on change over time

We found a strong temporal trend in both climate and plague risk since 1950 (Figure 3). To test whether this signal might be confounded by exogenous factors, we used a novel approach where we trained BART models with random intercepts (riBART) for each year, and projected the model again without random effects over the 68 year period (see Methods). The random intercepts identified interannual variation in prevalence (Extended Data Figures 13,14), detrended the data, and allowed the models to identify climate signal minus the confounder without substantially changing overall predictions (Extended Data Figure 15). Subsequently, we predicted how suitability changed using the same model without random intercepts; this allowed us to be confident that the changing suitability we identified in these “detection models” was the consequence of constant relationships between temperature, precipitation, and plague transmission.

**Figure 3:**
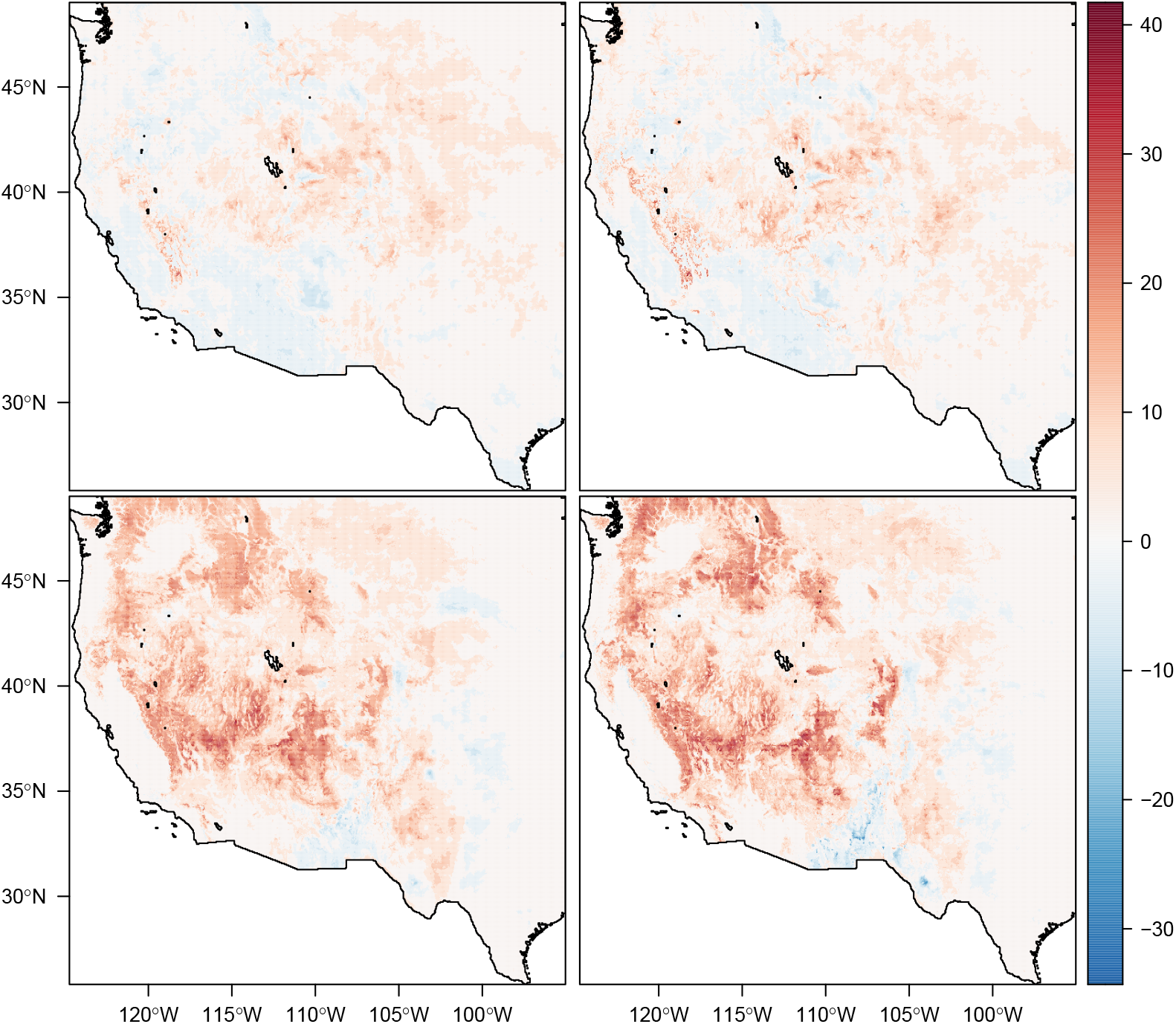
Total percent change in plague suitability, 1950 to present, in the human (top) and wildlife models (bottom), before (left) and after (right) adding random intercepts to control for interannual variation.

Both detection models identified a meaningful signal of temporal variation. The random intercepts identified a signal of rising prevalence through the wildlife dataset, particularly increasing after 2011 when the diagnostics were changed (Extended Data Figures 13,14). In the human model, we identified a much more subtle long-term quadratic trend peaking in the 1980s, matching a pattern that has been previously attributed to climate cycles like the Pacific Decadal Oscillation.^51^ Surprisingly, the “detection” models identified an even stronger pattern of change over time (Extended Data Figure 16). In the wildlife model, suitability increased an average of 4.8%, and 4.9% in the detection model, with a much fatter tail to the distribution as well. In the human model, suitability increased an average of 1.7% from 1950 to 2017, and 2.1% in the detection model. In much of the region, we found that plague suitability increased by 30 to 40% over the entire interval. We found that suitability rose most substantially in the wildlife model at high elevations, while spillover risk increased more gently, and peaked at mid-elevations (Figure 4, Extended Data Figure 17). Because the detection models only predict change across years based on temperature and precipitation, we are confident that the increase in the long term signal of warming (roughly 0.8° in the region since 1950) and higher anomalous precipitation are responsible for these predicted changes (see Extended Data Figures 18,19). We conclude that, even with several confounding factors, environmental change since the 1950s may have helped plague reservoirs become established at higher elevations, and slightly increased the risk of spillover into human populations at mid-elevations. Over the coming half-century, previous work suggests that the shift towards higher elevations is likely to continue in this region^54^.

**Figure 4:**
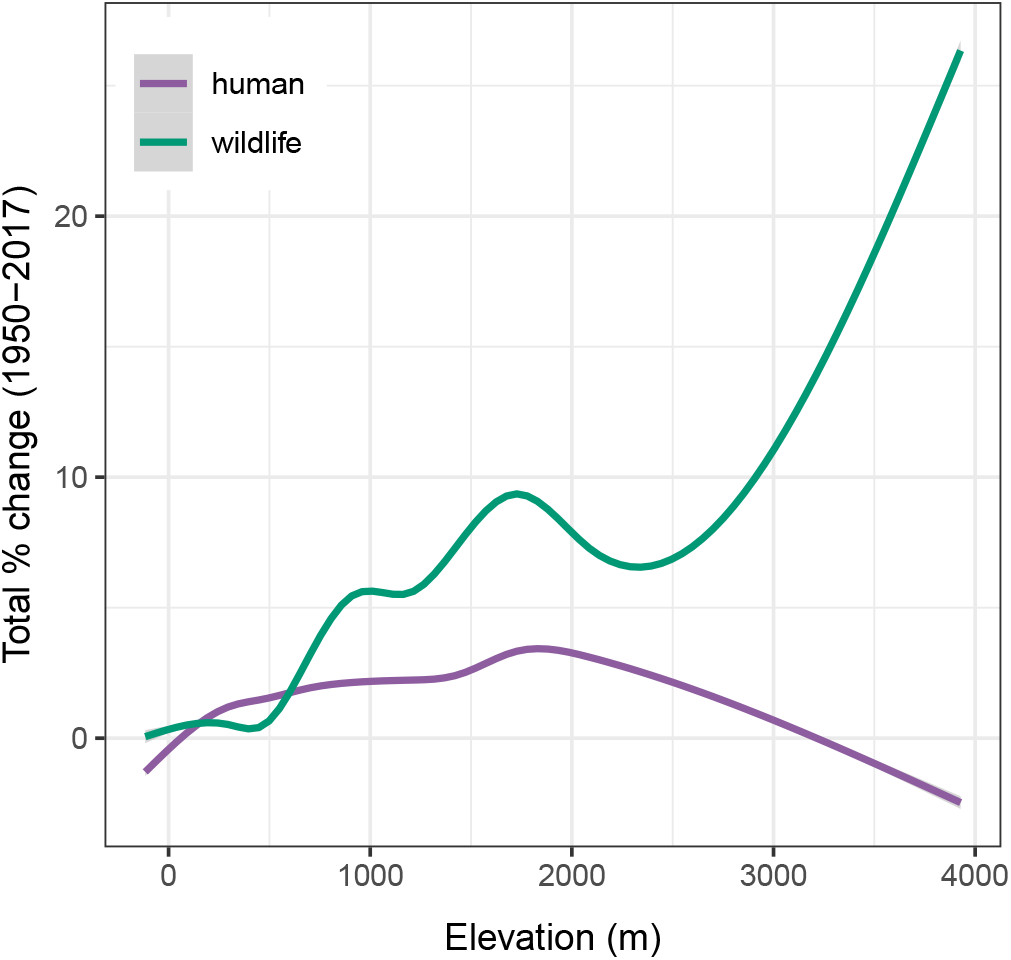
Environmental suitability for plague has increased substantially at high elevations for wildlife; risk of spillover has increased mildly at mid-elevations. Lines are given as generalized additive model smooth fits based on the detection models.

## Discussion

Our study shows that human and wildlife data can be used together to map plague reservoirs and spillover risk in the United States, and to make meaningful inferences about ecological drivers of transmission. We found support for two major hypotheses: the biodiversity amplification effect and the trophic cascade hypothesis. Support for these patterns has increasingly been found across systems, and points to a view of plague risk where weather conditions (and their impact on flea vectors) in rodent biodiversity hotspots are the primary driver of transmission and spillover. We further found strong evidence that the North American distribution of plague is heavily influenced by soil conditions. The global distributions of soil-persistent bacteria like anthrax (*Bacillus anthracis*), tularemia (*Francisella tularensis*), and botulism (*Clostridium botulinum*) are known to be constrained by the biochemical properties of soil. Less is known about plague, which is not spore-forming, and until recently was mostly thought to behave like a typical vector-borne zoonosis. It may be that soil properties affect the suitability of burrows for higher flea densities, or determine host homeostasis for minerals that impact the virulence of the infection; plague foci might therefore fall in the narrow range of conditions that can harbor higher densities of fleas, but do not substantially increase the lethality of the infection. However, increasing evidence also suggests that the bacterium can persist in the soil, possibly through symbiotic relationships with amoebas, for weeks to months—and possibly even years^37,35^. These complexities underscore the importance of a One Health approach while studying the ecology of plague, which—like anthrax and many other bacterial pathogens—circulates easily among fleas, rodents, other wildlife, humans, and the environment as one interconnected system^90^.

Developing a better understanding of plague in well-studied systems like the American West will help develop a broader picture of its ecology. At present, all global maps of plague foci have been compiled from expert knowledge; modeled products in the English language are limited to the western United States, China, and Africa (see Extended Data Table 2). In part, this reflects the challenges of sharing, aggregating, and consolidating surveillance data. It may also likely reflects concerns about model transferability, given that the complex multispecies dynamics of plague reservoirs differ greatly across ecosystems and continents. However, other pathogens with regional host communities and complex environmental persistence have been globally mapped through multinational coordination^38^, and the same synthesis is possible for plague. In the more immediate term, our model also strongly suggests that wildlife reservoirs extend up to both national borders, and could plausibly extend beyond them (recently confirmed for the northern border^91^), but the official World Health Organization map of plague (last updated 2016) includes no reservoirs in Mexico or Canada. Collaborating with national surveillance infrastructure in both countries may help resolve the boundaries of plague transmission more clearly, and reveal foci currently overlooked by global monitoring efforts.

Beyond plague, our study highlights the opportunity for medical geographers to develop new methods that are suited to a rapidly changing world. Here, we proposed two methodological advances that build on existing best practices in infectious disease mapping. First, time-specific covariates allowed us to train machine learning models on nearly a century’s worth of data, improving precision compared to coarsely-averaged predictors, and capturing the effects of environmental change. If this approach is integrated with others at finer temporal scales, such as those that consider seasonal aspects of transmission or spillover risk^19,20^, this could begin to set the foundation for an early warning system. Second, the use of random intercepts to remove data and detection biases, such as the serology method change in our data sample in 2011, is an important step towards testing climate change impacts using continuous-time data (similar to how econometric approaches resolve these problems in similar spatiotemporal analyses^7^). We propose that when this approach can be taken, it may be used as a first principles method for detecting the signal of environmental change in species’ habitats. This could be a particularly important step towards synthesizing the impacts of climate change on the shifting presence and absence of disease data, especially in cases where prevalence and incidence data are lacking and panel regression approaches cannot be applied. However, this work will still need to be followed by proper “attribution” work that compares predicted patterns to counterfactual scenarios without climate change; at present, all we can conclude with certainty is that weather conditions have changed in a way that trends favorably for plague risk.

Our study also points to a number of gaps in our understanding of environmental change (and consequently, potential methodological limitations). The PRISM data offers a fairly comprehensive view of the recent climate in the United States, and allowed us to identify the role of temperature and precipitation in plague transmission. However, we held both soil and rodent predictor variables constant, and neither are stationary in reality. Soil has changed over the last century due to a combination of climate change and land use change, and unfortunately time-specific covariates are unavailable; in many cases, our soil layers had to be generated custom to this study, and for the rest of the world these data are even more sparse. Similarly, evidence is strong that most terrestrial species have responded to recent climate change by undergoing range shifts, especially along elevational gradients. If rodents have undergone range shifts, they may have encountered novel vector communities, and the relationship between richness and transmission could change. Similarly, if elevation acts in our models as a proxy for specific rodent-flea assemblages, range shifts could decouple the observed relationships between elevation and transmission. As other studies have pointed out, these challenges highlight the need to begin integrating zoonotic surveillance and biodiversity monitoring^1^.

## Methods

Despite recent interest in modeling the distribution of major infectious diseases^92,10^, there is no definitive global map of plague reservoirs. All existing global plague maps have been derived from expert opinion^93^; all modeled products so far have been produced for national or continental scales (see Extended Data Table 2). Plague ecology is regionally variable enough that this patchwork approach has the advantage of being tailored to relevant local predictors. However, the mix of modeling methods, variables, and spatiotemporal scales makes it nearly impossible to compare these models and develop any consensus on the biological or geological factors that determine where plague reservoirs can exist, and where not. In this study, we adapt predictors that have previously worked in other similar work on plague, and develop novel models of spatiotemporal risk patterns in the western United States based on human and wildlife data spanning 1950 to 2017.

### Data

Our study is designed around two independently-collected datasets, with only a small amount of temporal overlap. Together, they provide as comprehensive a picture of plague in the United States as possible.

#### Human case data (1950-2005)

Human cases of plague occur sporadically but consistently in the Western United States, driven partially by exposure to infected cats and dogs that have acquired the infection outside of the home. The vast majority of cases are bubonic, though a handful of pneumonic and septicemic cases occur. Confirmed plague cases are mandatorily reported to the U.S. Centers for Disease Control and Prevention (CDC) Emergency Operations Center. CDC surveillance data is actively maintained on plague, and has been previously published in summary form as county totals^51^. We re-used these data, which have been anonymized by previous researchers, and had case geolocations aggregated to county totals. To georeference them, we randomly sample a number of locations within each county equivalent to annual case totals. A total of 860 plague cases are recorded over the interval, with an average of 7.7 cases per year, across 490 counties in the American west.

#### Wildlife serology data (2000-2017)

Wild animals are routinely exposed to *Y. pestis* in endemic regions, including the United States. Infection leads to substantial morbidity and mortality in some species (e.g., prairie dogs), but other species (e.g., coyotes) readily survive infection, with antibodies to *Y. pestis* being the only indication of exposure. This is especially true for predators, which can be exposed through consumption of plague-positive rodents or through bites from plague-positive fleas. These predator species do not necessarily play a direct role in plague transmission and dynamics, but instead act as sentinels of plague activity on the landscape^94,95^. Correspondingly, the USDA National Wildlife Disease Program tests wildlife for evidence of plague exposure throughout much of the western U.S. Testing was conducted using a hemagglutination assay^96^ at the Centers for Disease Control and Prevention until 2011. A majority of samples collected after 2011 were tested using a bead-based flow cytometric assay with a lower limit of detection^97^. In total, the version of the dataset we used spanned February 13, 2000 to January 29, 2018, with a total of 41,010 records, including 5,043 animals that tested positive. Of those records, the vast majority are coyotes (32,825 animals including 4,812 that tested positive).

### Environmental covariates

The transmission ecology of plague shares features with both vector-borne systems (e.g., malaria or dengue fever) and soil-borne pathogens (e.g., anthrax or melioidosis). The predictors we have chosen here are informed by predictors that have performed well for other plague mapping projects (see Extended Data Table 2), and were all expected to be informative as drivers of host ecology, vector competence and/or soil persistence.

Most studies that map infectious diseases with machine learning methods (i.e. ecological niche models) use long-term climate averages, paired with occurrence data that sometimes span decades of unstable environmental conditions. In contrast, we used time-specific climate data paired with—and extracted for—the year of each data point in the occurrence data. This allowed us to make yearly spatial predictions of the distribution of plague risk, and consider the extent of transmission risk as a dynamic process rather than a static surface. We held non-climate predictors constant, assuming them to either be invariant (elevation) or long term averages (soil and rodent richness); in a more advanced retrospective, it might be possible to reconstruct the impacts of land use change by adding yearly resolution to these covariates, but these data do not currently exist.

#### Climate

We derived all climate data (1950–2017) from PRISM, a historical reconstruction of climate in the continental United States, derived from a mix of weather station data and climatologically-aided interpolation.^98^ From the PRISM dataset, we used cumulative annual precipitation and annual mean, minimum, and maximum temperatures. We also generated two “anomaly” variables, given on a pixel-by-pixel basis as the difference between the annual value and the long-term average, divided by the variance. These data were downloaded in 2.5 arcminute grids (~4.5 km at the equator), which was used as the standard resolution for the rest of the project.

#### Soil

We assembled a set of seven predictor layers for soil persistence of plague that were informed by both laboratory experiments on plague transmission, and previous efforts mapping soil-borne pathogens like anthrax (*Bacillus anthracis*) and botulism (*Clostridium botulinum*). We aimed to develop a cohesive set of predictors characterizing the C layer (~ 1m depth); rodent burrows in the American west can go up to two meters deep in the soil, but macronutrient data is limited at this depth. We used the most recent version of the ISRIC SoilGrids global dataset at 250 meter resolution^99^, and selected gridded layers of soil pH, cation exchange capacity (base saturation), and the concentration of sand, clay, and organic content in the top 60-100 mm layer of soil. Sodium, calcium, and iron concentrations were derived from a national survey of soil geochemical properties, published in raw form as USGS data series 801.^100,101^ We extracted all point samples of mineral concentration in the C horizon, given in weight percent, and then developed a rasterized layer for these macronutrients by kriging the point data, using the autoKrige function in the automap package.

#### Additional covariates

Rodent species richness was derived by stacking species IUCN expert range maps for the Rodentia, and rasterizing the richness layer using the `fasterize` package. Elevation data was scraped using the `elevatr` package in R, which pulls gridded elevation data from the AWS Open Data Terrain Tiles. We pulled elevation data at resolution “6”, which returns elevation rasters in 2,446 meter squared grids at the equator (~1.3 arcminutes), and aggregated to the native resolution of the other grids.

### Modeling

Dozens of statistical methods have been applied to species distribution modeling in the past few decades, with a wide range of performance.^102^ Over the past few years, classification and regression tree methods (CART) – including random forests and boosted regression trees – have become especially popular for mapping the geographic distribution of infectious diseases^103,38,104,105,106,107^. Here, we use a fairly new method, Bayesian additive regression trees (BARTs), implemented with the `R` package `embarcadero` as a species distribution modeling wrapper for the dbarts package.^53^ BART is a powerful new method with growing application in computer science, and often performs comparably to other CART methods like random forests and boosted regression trees.^108^ In the `embarcadero` implementation, BARTs have several unique features that make them a powerful tool for disease mapping, such as: model-free variable importance measures, and automated variable selection; posterior distributions on predictions, as a measure of uncertainty; posterior distributions on partial dependence plots; two-dimensional and spatially-projected partial dependence plots; and various extensions, including random intercept models.

Like other CART methods, BART makes predictions by splitting predictor variables with a set of nested decision rules (“trees”) that assign estimated values to terminal nodes (“leaves”). Predictions are generated based on a sum-of-trees model, where a set of *n* trees with leaves (*T*_1_, *M*_1_),…, (*T_n_, M_n_*) each make predictions *g*(·) that are added together, for a total estimate:

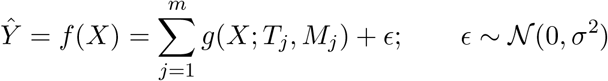

For logistic classification problems (like species distribution modeling), BART uses a logit link function:

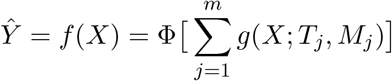

where Φ is the standard normal cumulative distribution. An initial set of *n* trees is fit, and then altered in an MCMC process based on a set of random changes to the sum-of-trees model (e.g., new splits added, levels rearranged, or leaves pruned). An initial burn-in period is discarded, and then a set of posterior draws of *f* ^∗^ create the posterior distribution for *p*(*f*|*y*) ≡ *p*(trees|data).

BART is easily implemented out-of-the-box, even with a full Bayesian MCMC component. Three priors control the ways decision trees change: the probability each variable is drawn for a split, the probability of splitting values tested, and the probability a tree stops at a certain depth. In the simplest form, the first two can be set as uniform distributions, while the latter is usually set as a negative power distribution; they can also be adjusted using a full cross-validation approach. This is handled automatically in the `dbarts` package, for which `embarcadero` is a wrapper. More advanced implementations with complex prior design are sometimes appropriate: for example, a Dirichlet distribution on the variable importance prior can help identify informative predictors in high dimensionality datasets (dozens or hundreds of covariates). However, in our case, we had confidence all variables were biologically plausible based on expert opinion.

#### The base models

We ran two separate baseline models, the first using human data from 1950 to 2005, and the second using the wildlife data from 2000 to 2017. For the human model, we used the number of cases recorded each year in each county to generate a set of random georeferenced pseudopresence points. We then generated seven pseudoabsence points in each year to create a roughly balanced design, for a total of *n* = 430 pseudopresence points and *n* = 392 pseudoabsence points. For the wildlife model, we balanced the design by subsampling seronegative animals in equal number to seropositive ones, for a final *n* = 5, 002 true presence points and *n* = 4, 759 true absence points.

Both models were run with the full predictor set, followed an automated variable set reduction procedure implemented in embarcadero that formalizes the recommendations of Chipman *et al.* (2010).^108^ In BART, variable importance is “model-free,” measured as the number of splitting rules involving a given variable (but incorporating no information on the proportional effect on the outcome variable, or proportional improvement of the model predictions). In models with fewer trees (small *n*), informative variables tend to be selected more often, while uninformative variables are selected rarely or drop out entirely. This property of BART establishes a rubric that can be used to identify an informative variable set. First, an initial model is fit with all variables 100 times each for six different settings of ensemble size (*n* = 10, 20, 50, 100, 150, and 200 trees). Plotting the average importance of variables at each level offers a qualitative diagnostic of how informative each predictor is. Next, an initial set of 200 models with *n* = 10 trees are run, and variable importance is recorded and averaged across models. Models are run again (200 times) without the least informative variable from the first fit, and this is performed iteratively until only three variables remain; the variable set with the lowest average model root mean square error (RMSE), and therefore highest accuracy on the training data, is selected. Finally, we plot variable importances (including standard deviations based on model permutations).

Final models were run with the reduced variable set, with recommended BART model settings (200 trees, 1000 posterior draws with a burn-in of 100 draws) and hyperparameters (power = 2.0, base = 0.95 for the tree regularization prior, which limits tree depth). We then used the `retune` function in `embarcadero` to run a full cross-validation panel on the three prior parameters. `retune` runs a full cross-validation across the k hyperprior (values of 1, 2, and 3), the base parameter (0.75 to 0.95 in increments of 0.05), and the exponent parameter (1.5 to 2 in increments of 0.1), and returns the model with the parameter combination that generates the minimum root mean squared error.

For the wildlife model, the final variable set included: temperature mean, maximum, and anomaly; rodent richness; elevation; and five soil traits (calcium, sodium, iron, clay, and sand). The model validated well on training data (AUC = 0.836). For the human model, the final variable set included a similar subset: precipitation anomaly; temperature mean, maximum, and anomaly; rodent richness; elevation; and four soil traits (sodium, iron, clay, and calcium). The model also validated well on training data (AUC = 0.909).

#### Alternate formulations

As a final check of model performance, we ran a separate model with the same predictor sets that withheld the years 2000–2005 from both. On the test dataset for humans (n = 64), the model performed very well by the standards of external cross-validation (AUC = 0.820); on the test data for wildlife (n = 796), the model also performed well (AUC = 0.775). This indicated that both models were performing adequately.

We also recognize that model design can have a substantial effect on machine learning performance, and the downstream biological inference made by using ecological niche models. Given that BART is a relatively new method, it has been comparatively underexplored in this regard, and so a standard panel of “best practices” has not yet been recommended in the literature. However, for transparency about model uncertainty and the influence of subjective decisions on model outputs, we produced four major alternate formulations. First, we produced models that included all variables, rather than using the variable set reduction procedure, for both the human data (Extended Data Figures 20,21, and wildlife data (Extended Data Figures 23,24, 25). We additionally considered two alternate formulations of the wildlife model. In the first, we used pseudoabsences instead of the true absences available in the data (Extended Data Figure 26). Though this increased model AUC (0.929), and allowed slightly different balancing of the data, it lead to visually-apparent overfitting. Finally, we ran an alternate model only using the coyote data in the NWDP dataset, which also performed adequately (AUC = 0.826; Extended Data Figure 27). Both models were ultimately not selected because they left available, biologically-meaningful data unused, and both produced predictions that were slightly less congruous with the human model.

#### Prediction, delineating foci, and measuring change

Although the models were trained over different intervals, the continuous and standardized set of predictors allowed cross-prediction over the entire extent of the study (1950–2017). For each layer of annual prediction, we thresholded suitability based on a model-specific threshold chosen to maximize the true skill statistic on the test data. We mapped areas of “unstable foci” as any region with at least one year of suitability, and “stable foci” as any region suitable in every year over the 70-year interval. This allowed us to compare long-term spatial patterns between the two models.

#### Random effects models for interannual variation

Prevalence changes year-to-year in both the data and modeled landscapes, but detecting the signal of climate change in that fluctuation can be challenging. There are several reasons prevalence could vary across years: (1) incidence is stochastic but temporally autocorrelated; (2) normal climatic variability (e.g. the Pacific Decadal Oscillation) or other socioecological trends (e.g., rising human populations) might also contribute to interannual variation, including non-linear trends over time; (3) anthropogenic climate change is directly driving changes in plague risk, or indirectly changing the ecology of the involved species; (4) sampling effort varies between years (for wildlife); or (5) detection rates could change between years, due to testing or surveillance. The last of these is particularly relevant as a possible confounder, given that wildlife diagnostics changed in 2011. A positive trend in plague risk might be generated by increased climatic suitability for plague tranmission, but could also be generated by a consistent increase in plague detection due to improved diagnostics and increased sampling effort, loosely colinear with warming temperatures on the scale of 20 to 70 years.

We propose a new method that uses machine learning approaches (i.e., ecological niche models) to detect the signal of environmental change while adjusting for confounders at a high level. The approach is loosely modeled off the ideas underlying econometric approaches to climate change detection and attribution, which usually use fixed effects panel regression to control for spatiotemporal confounders in climatic signal. By attributing as much variance as possible to spatial, temporal, and other confounders, and then identifying climatic signal in the remaining variance, these approaches can pinpoint the signal of environmental change with a high degree of confidence. So far, no analog to these approaches exists for ecological niche models. Only a handful of studies have even added temporal heterogeneity to ENMs; so far, we know of none that have also independently controlled for interannual variation in detection, sampling effort, or species prevalence.

A solution to temporal confounders is particularly needed in this study, given the challenges of the time-specific approach. In default settings, BART predictions converge on observed prevalence, i.e., *Ŷ* = *E*[*Y*] = *P* (*Y* = 1). For this reason, we balance the presences and absences, so that the model is as close to 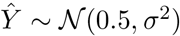 as possible. This produces a unique challenge for time-dependent modeling. Presences are distributed unevenly across years, and consequently, so is positivity. In the human model, this arises artificially, because pseudoabsences are generated evenly across years. We chose this approach to avoid over-representing years with more cases in the data, which would introduce an additional colinearity, but as a result relative prevalence varies substantially. This bias also affects the wildlife model more organically; although the number of points per year varies independently of test positivity (cor = 0.113, *p* = 0.687), because most sampling is passive, there is still a wide range in annual prevalence (28% in 2006 versus 80% in 2017, both years with several hundred records), with a net trend towards higher positivity over time. Because prevalence varies between years in both models, the resulting colinearity with environmental change could confound the detection of meaningful signals.

Inspired by the econometric approach, we propose a use case for the *random intercept BART model* (riBART), which has recently been proposed as an extension of the method for clustered outcomes. The approach adds a random intercept term to the model (separet from the tree-fitting process) based on the identified *K* clusters

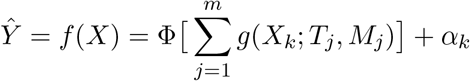

where the random intercepts *α_k_* (*k* ∈ 1: *K*) are normally distributed around zero (i.e., the *K* groups are assumed to have normally distributed, independent additive effects on the outcome variable). The error structure of the random effects and the sum-of-trees model are assumed independent. Here, we propose that the model can be fit as usual with a random intercept for year, as a way of accounting for temporal heterogeneity as a possible confounder. The yearly random intercept absorbs most of the interannual variation in plague prevalence (i.e., the relative ratio of presences and absences in the data), such that residual variation in prevalence should be roughly constant across years. Identifying the climatic signal in this residual data, and then examining predicted prevalence (without random intercepts) based only on environmental change, allows more confident statements about how environmental change contributes to shifting disease risk.

We revisited the two main models, and used riBART to add an annual random intercept to our model for each year, which we refer to throughout as the “detection” models. Fitting climate-plague response curves after this detrending decouples the possible colinearity between climate trends and coarse interannual signal in the data, which may be driven by natural variation in prevalence or other confounders (e.g, the 2011 change in wildlife testing protocols). We fit both detection models with a random intercept for year, plotted the random effects, and predicted over the 70 year interval without the random effect included. (All functionality to implement SDMs with riBART is available in `embarcadero` as an updated release.)

#### Detecting change over time

To estimate trends of change over time, we fit a linear slope through each pixel-by-year. Multiplying by 68 years, we were able to estimate total percent change in suitability since 1950 in a given pixel. We did not limit these to pixels with a significant trend, as any frequentist significance test iterated over millions of pixels would be mostly meaningless. We generated these maps for the two primary models and the two detection models (Figure 3), as well as (in the supplement) for mean temperature and precipitation (Extended Data Figures 18,19).

### Data and code availability

Human case data in this study is taken from previous studies and is available online for researchers to reproduce our study. Wildlife data is available on formal request and approval from the United States Department of Agriculture. All code is available at github.com/cjcarlson/plague-wna.

## Supporting Information

**Extended Data Figure 1:**
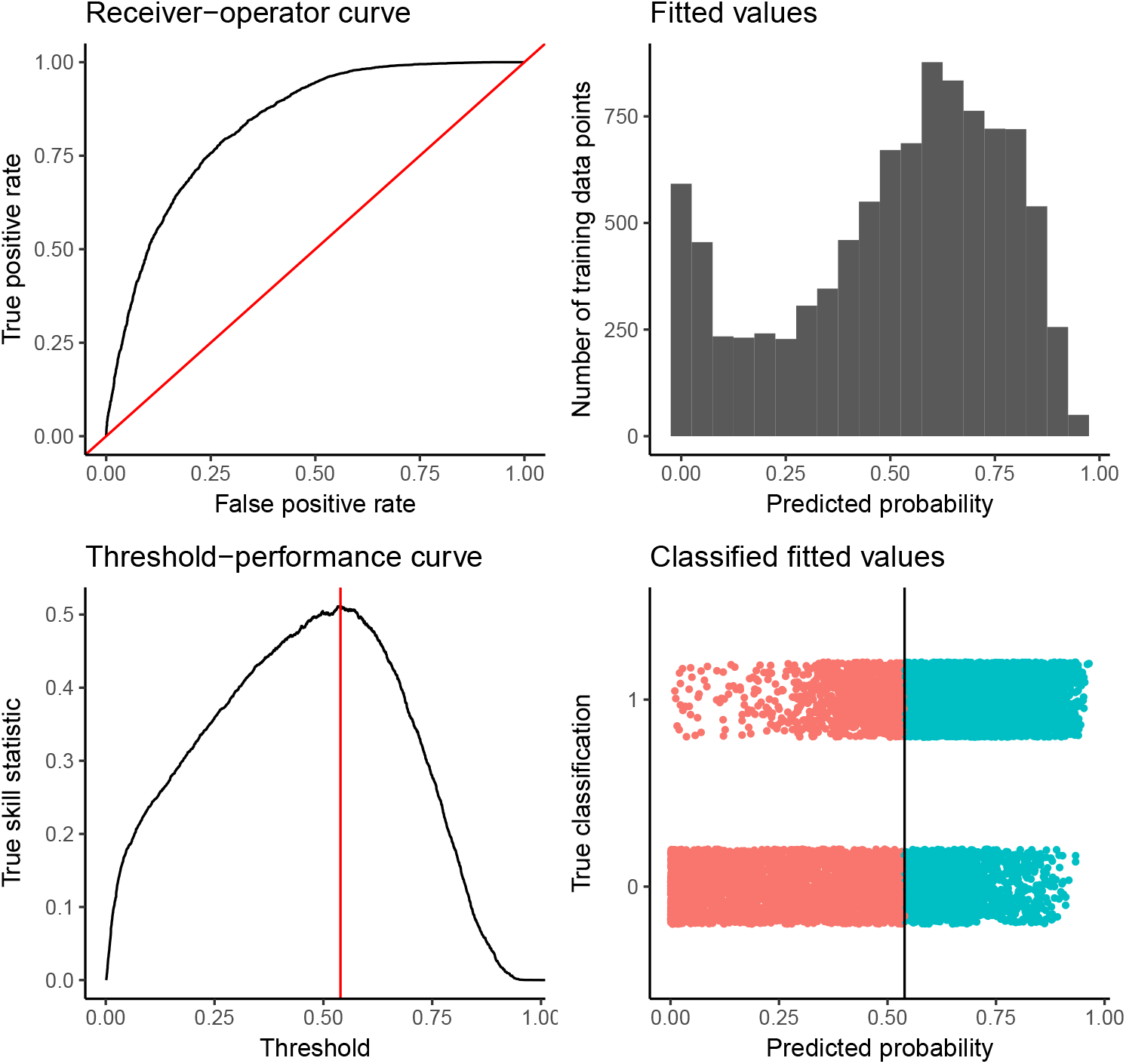
Summary model diagnostics for the wildlife plague risk model.

**Extended Data Figure 2:**
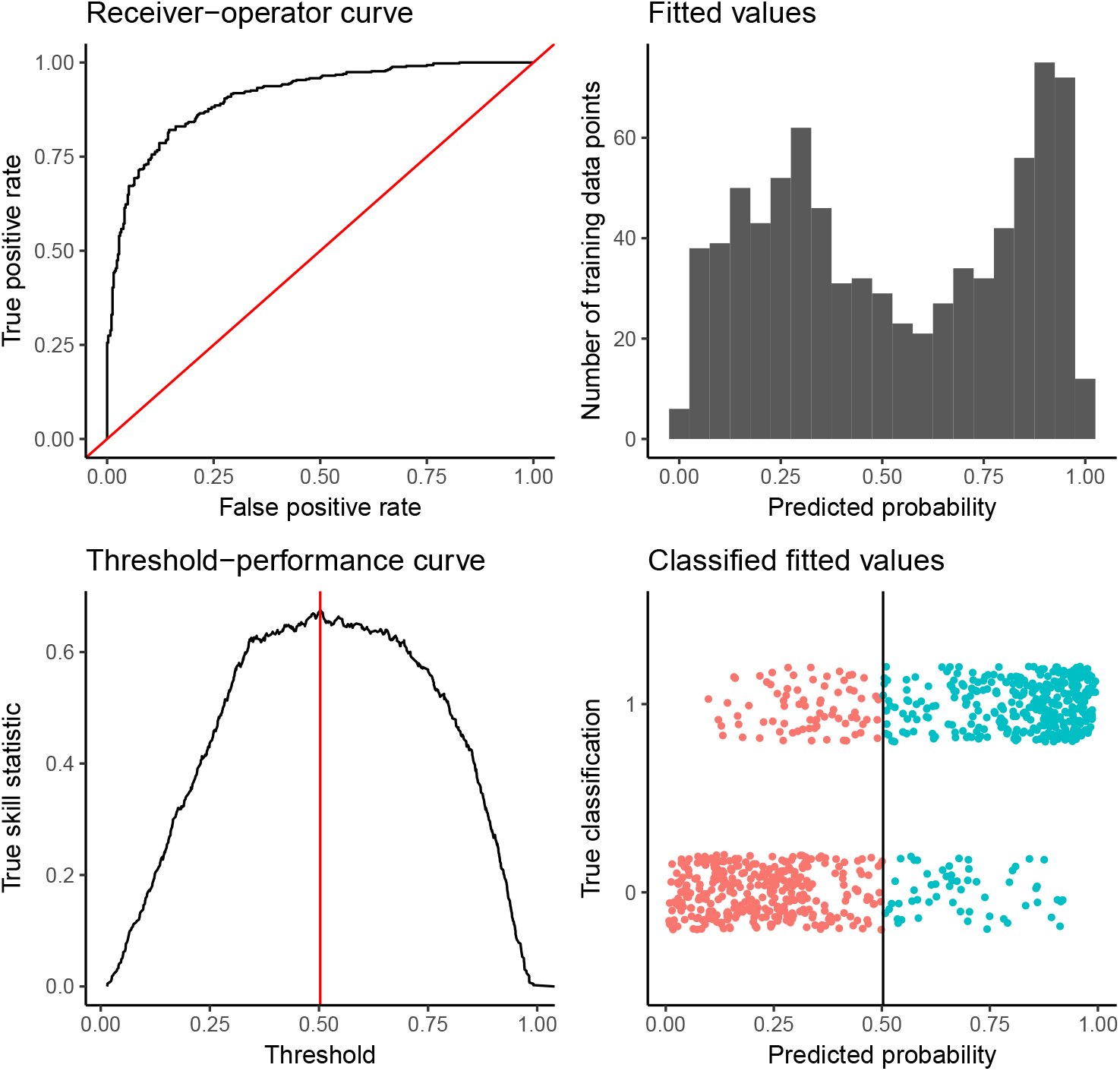
Summary model diagnostics for the human plague risk model.

**Extended Data Figure 3:**
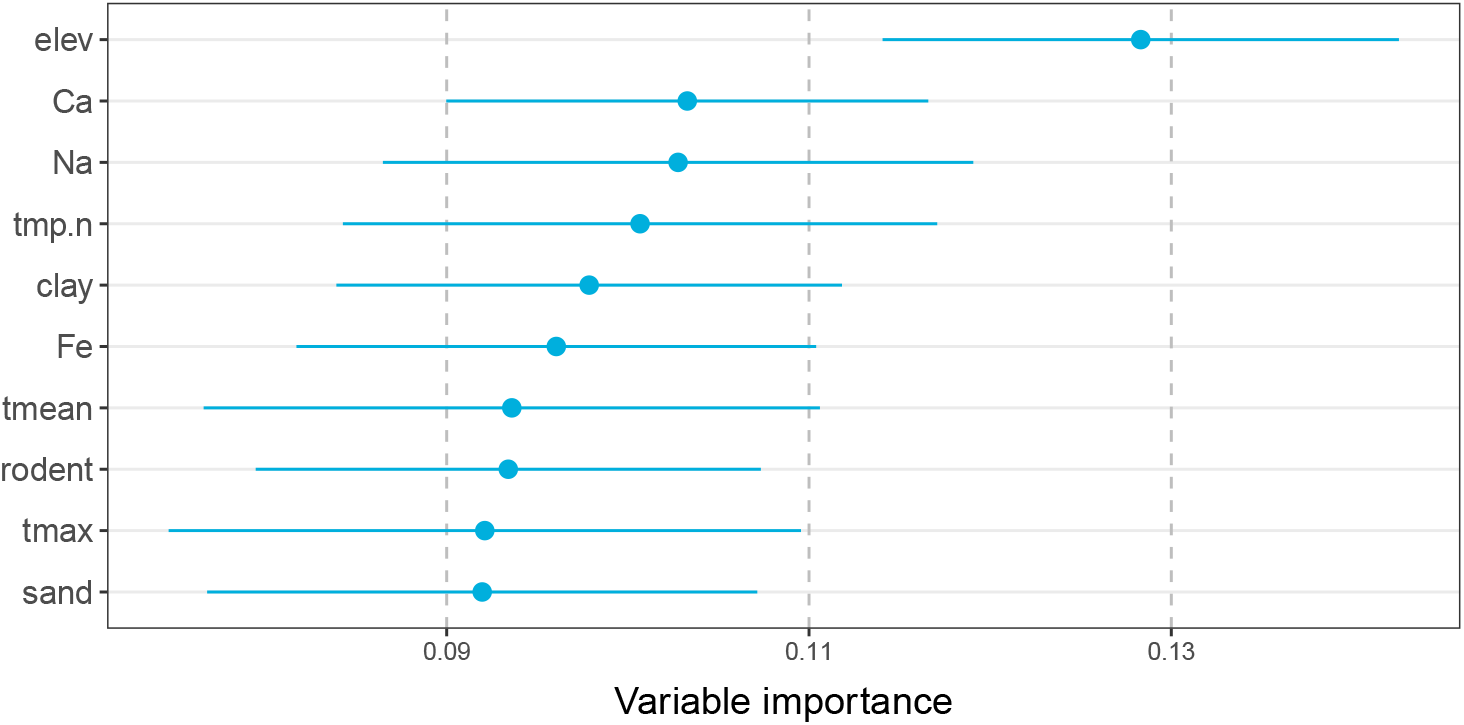
Variable importance in the wildlife plague risk model.

**Extended Data Figure 4:**
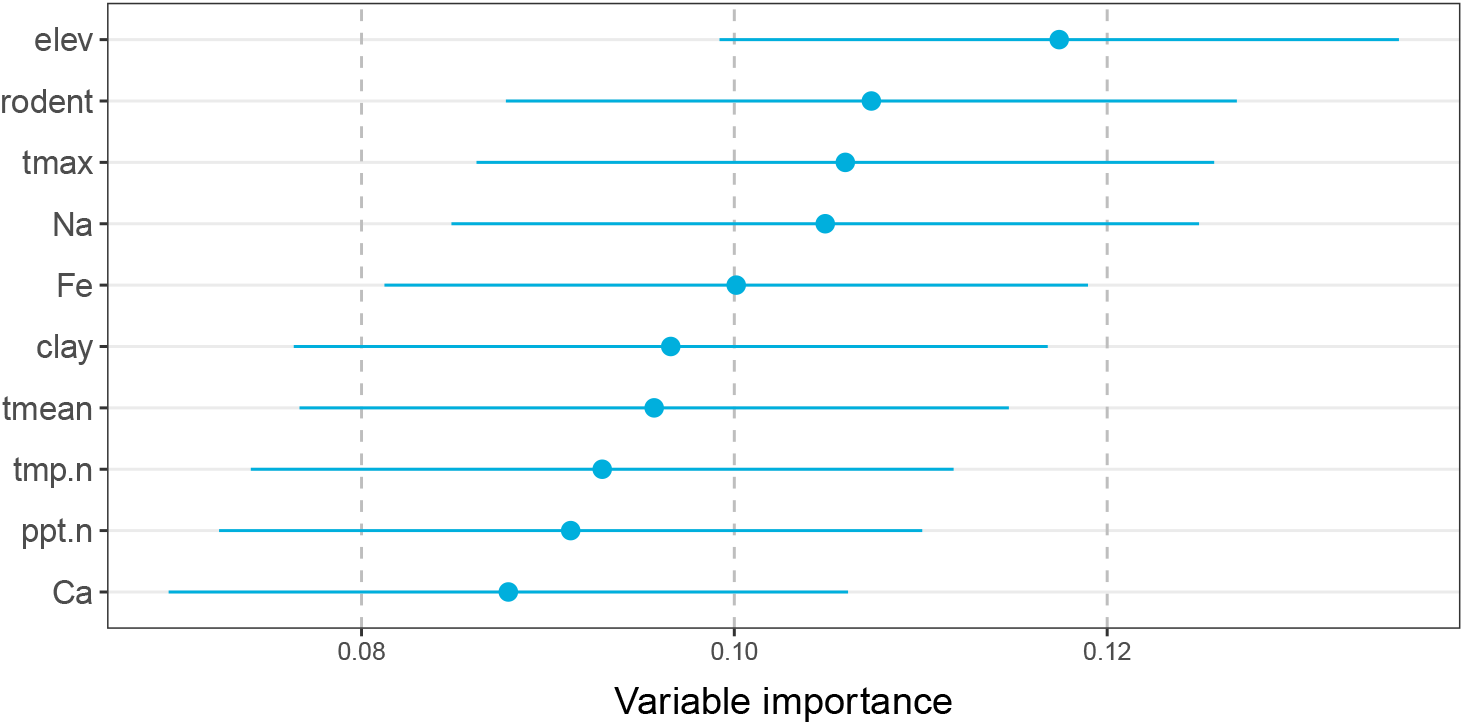
Variable importance in the human plague risk model.

**Extended Data Figure 5:**
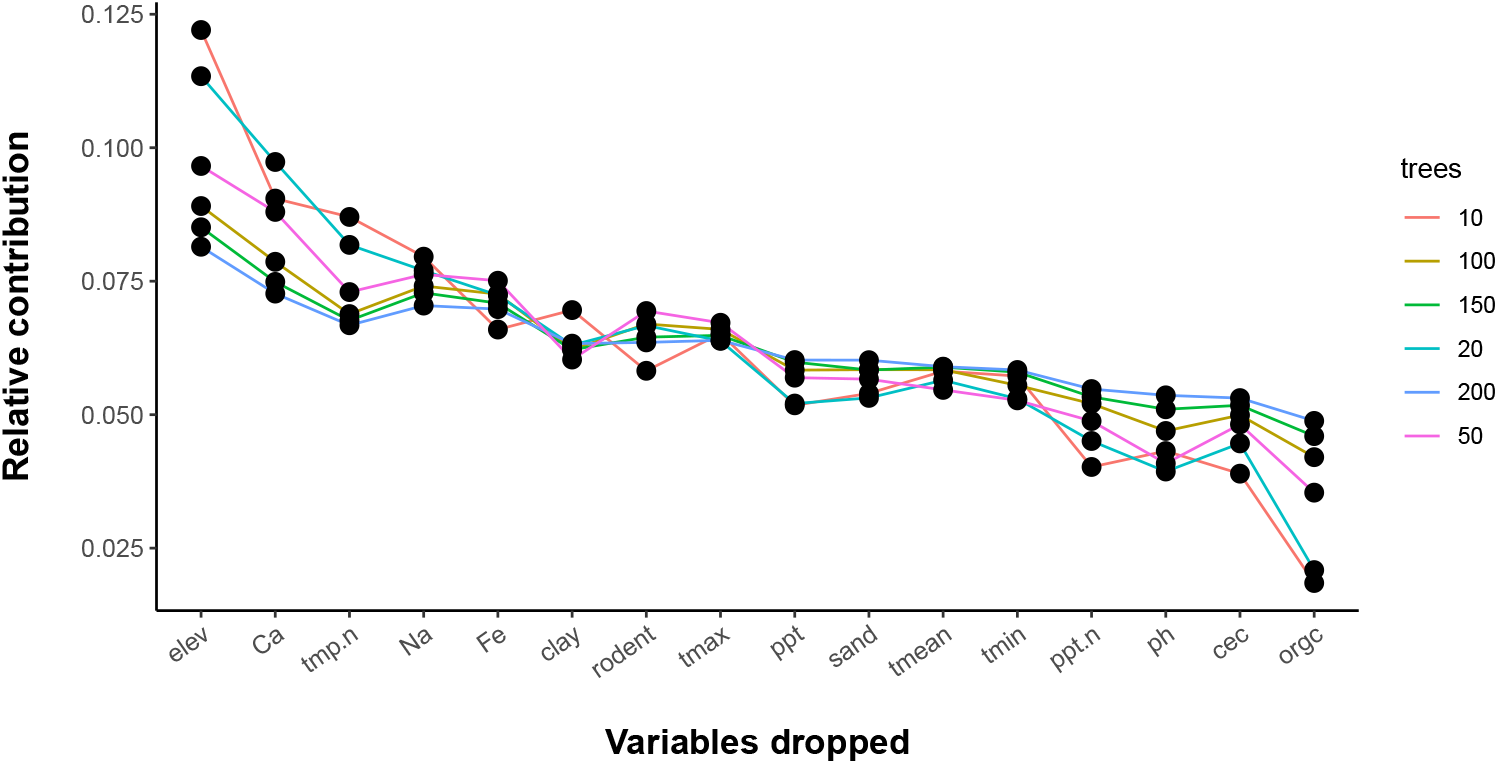
The variable importance diagnostic for all variables considered for the wildlife plague risk model.

**Extended Data Figure 6:**
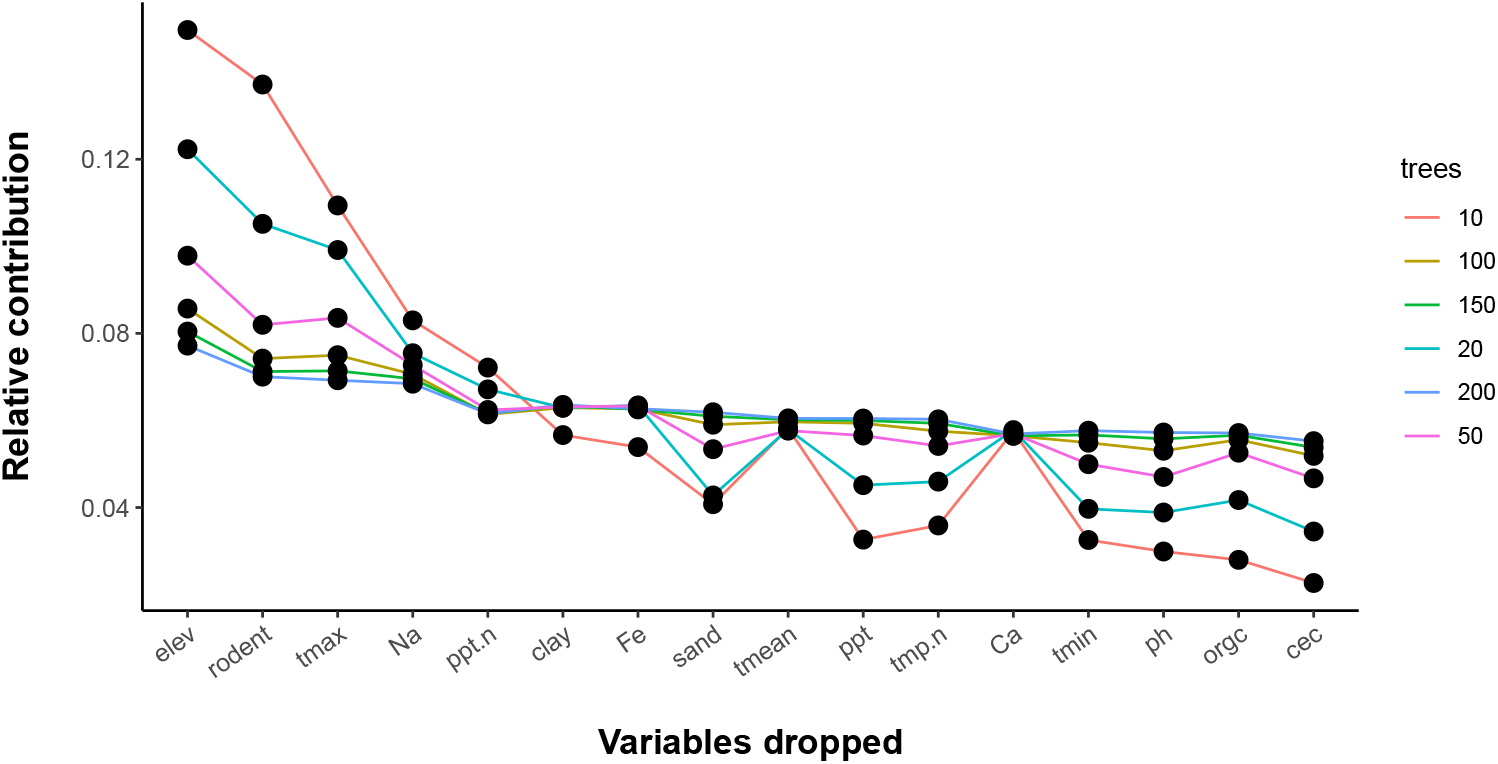
The variable importance diagnostic for all variables considered for the human plague risk model.

**Extended Data Figure 7:**
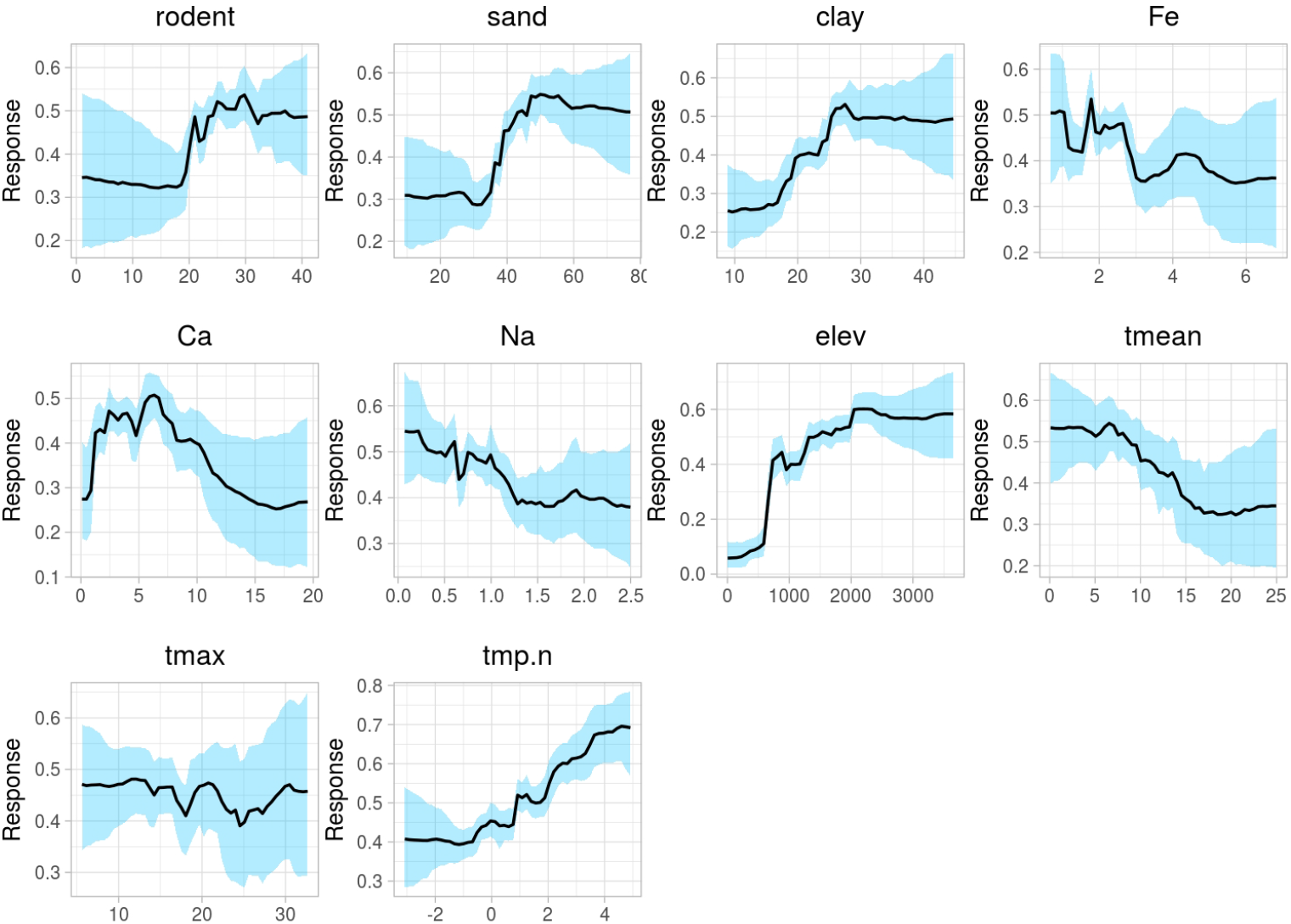
Full partial dependence plots for the wildlife plague risk model.

**Extended Data Figure 8:**
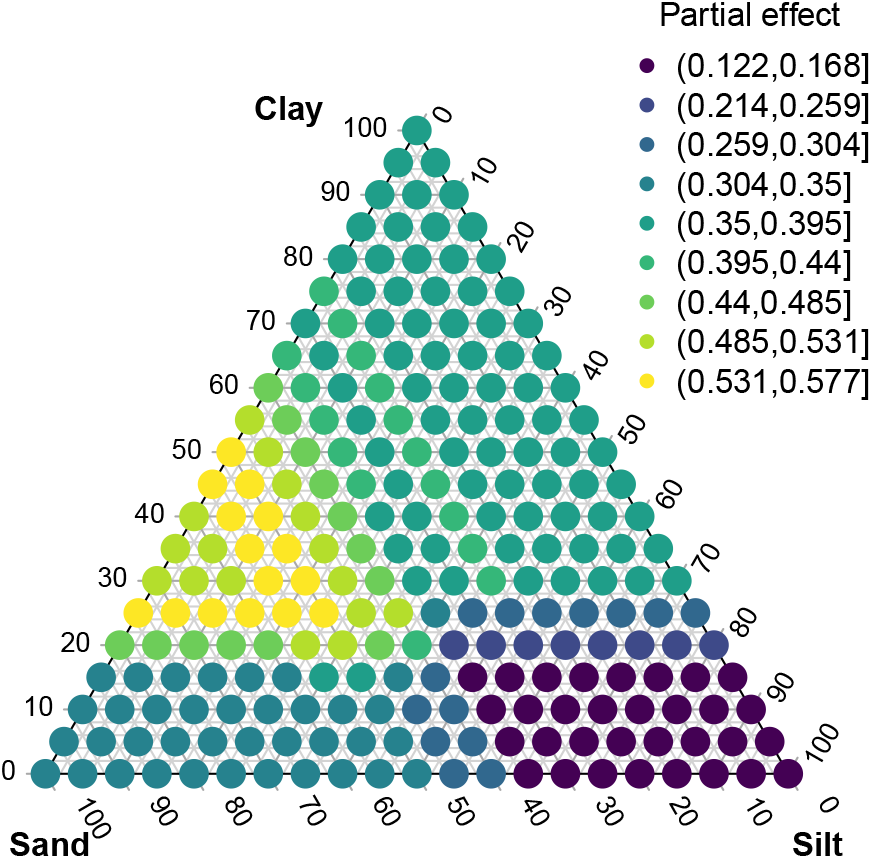
Two-dimensional partial dependence plot for sand and clay in the wildlife model, projected onto the soil composition triangle as a ternary partial (tertial) plot.

**Extended Data Figure 9:**
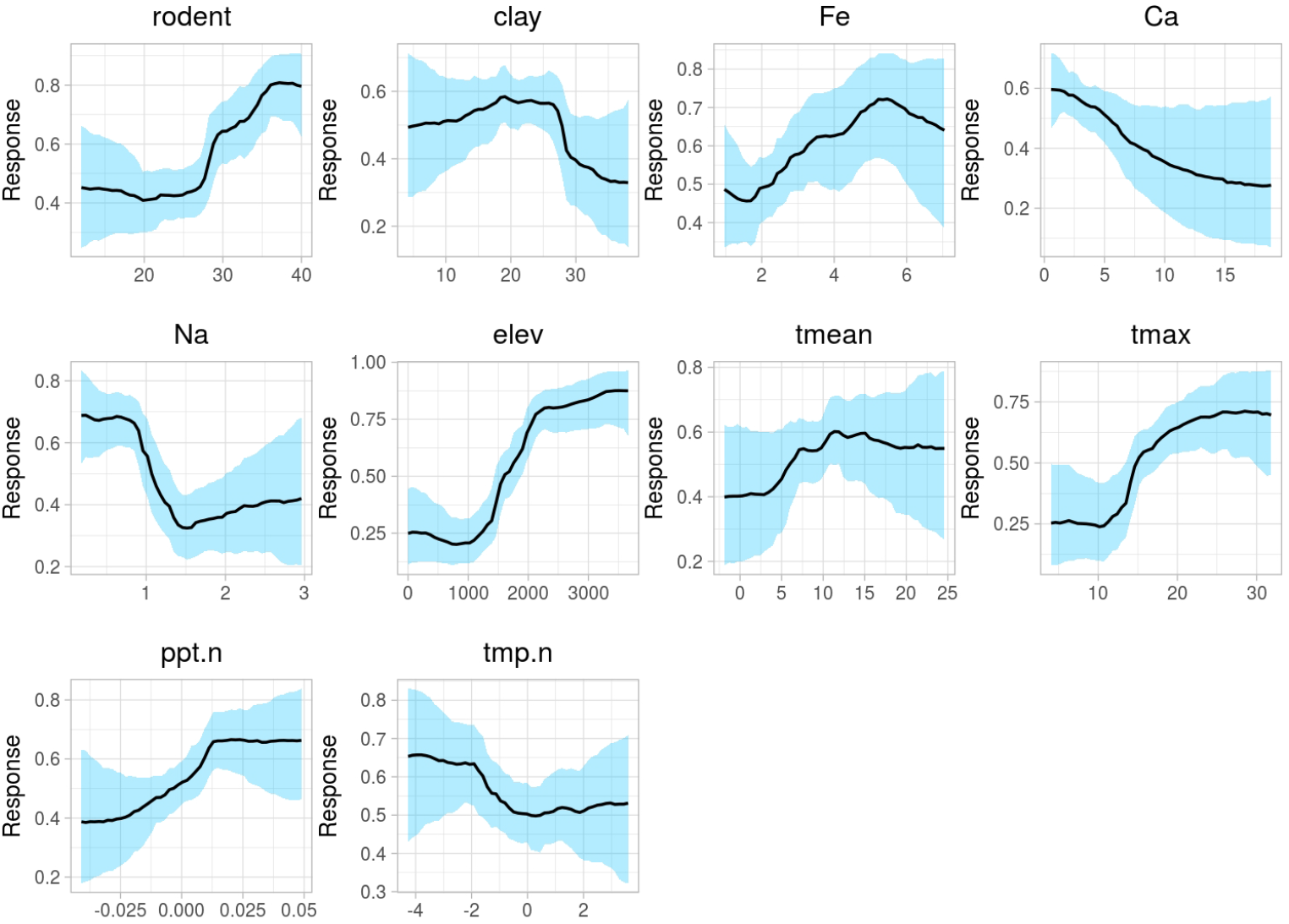
Full partial dependence plots for the human plague risk model.

**Extended Data Figure 10:**
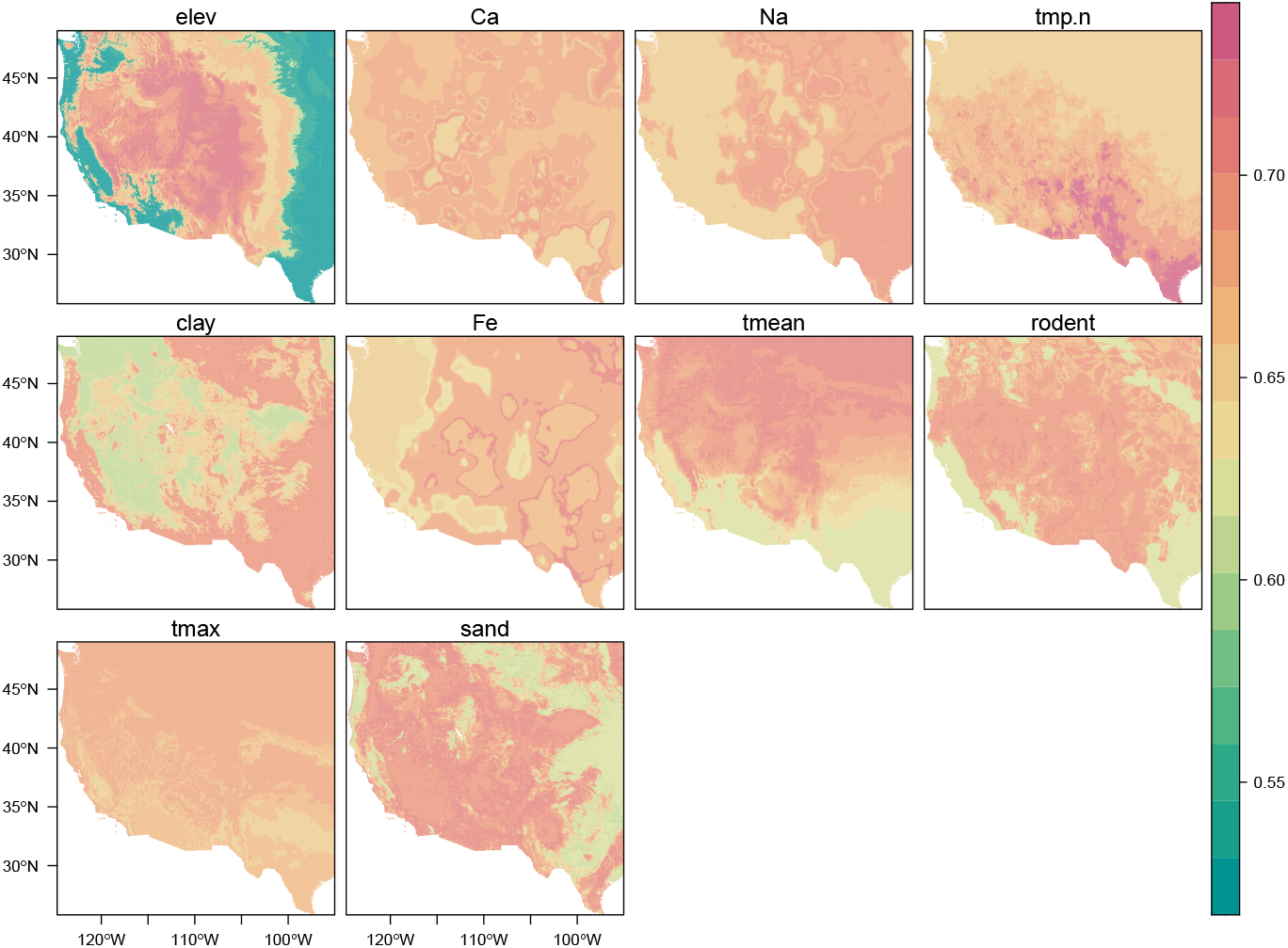
Spatial partial (spartial) dependence plots for the wildlife plague risk model. Variables are given in descending order of model importance.

**Extended Data Figure 11:**
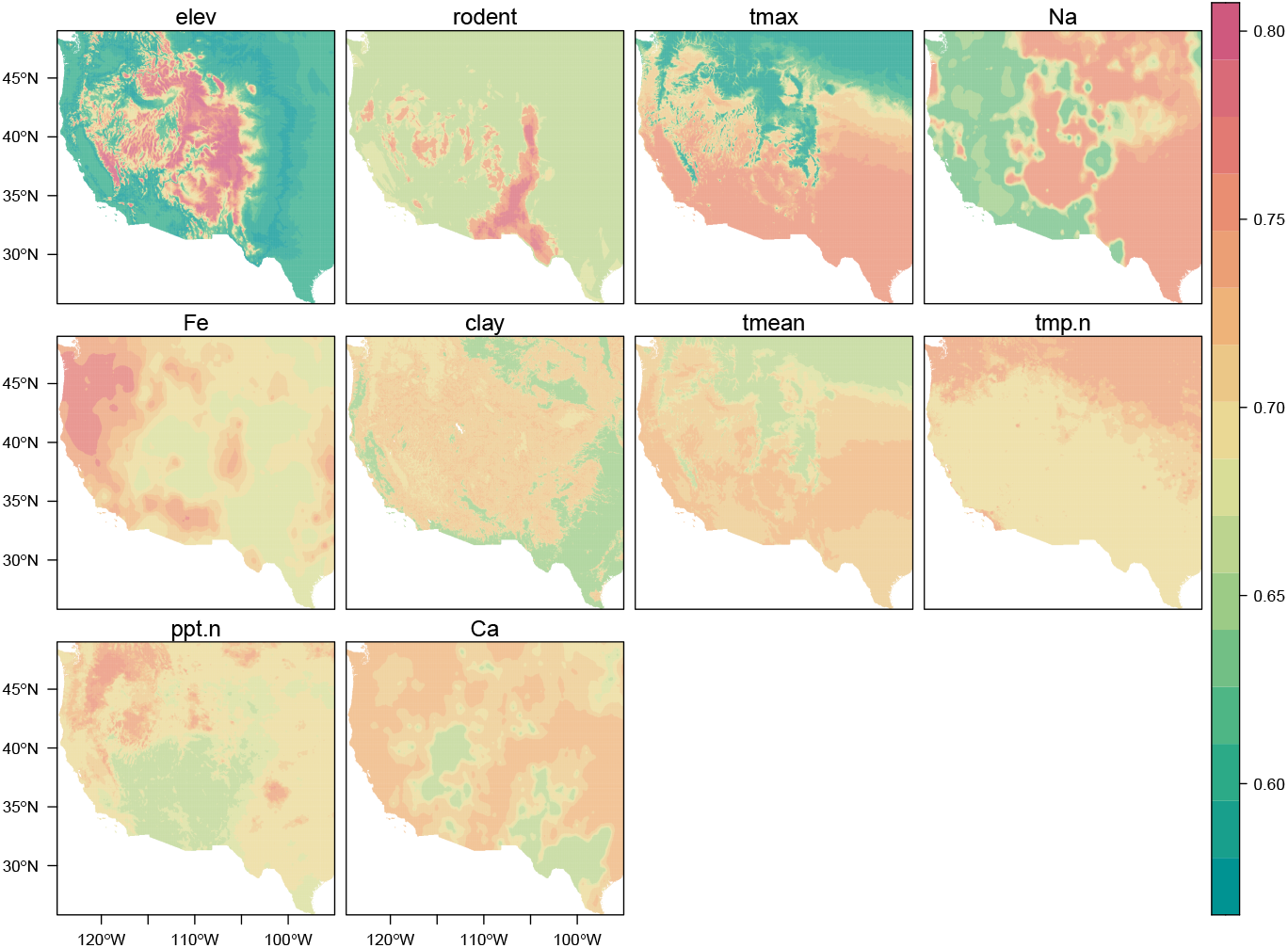
Spatial partial (spartial) dependence plots for the human plague risk model. Variables are given in descending order of model importance.

**Extended Data Figure 12:**
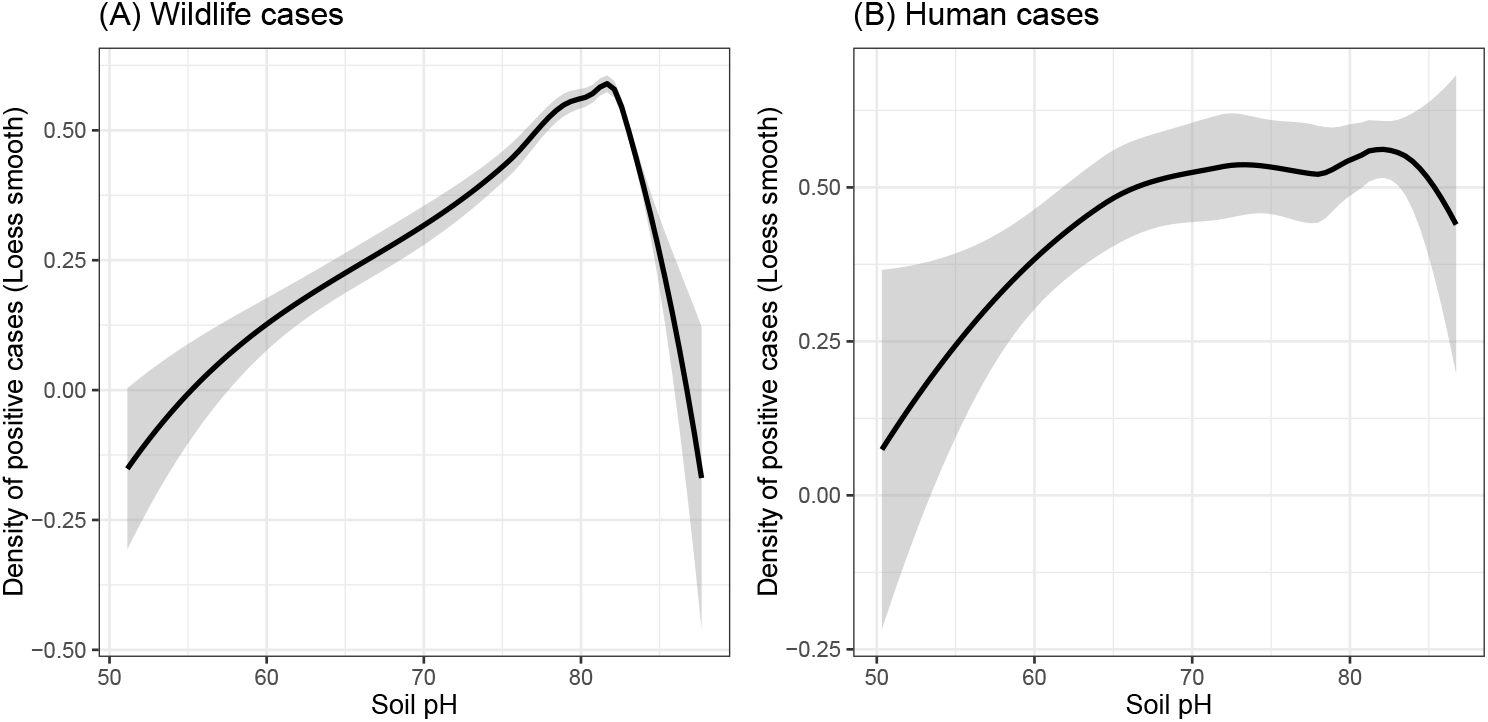
Density of positive cases in the wildlife and human case data relative to recorded pH, modeled using a simple Loess smooth.

**Extended Data Figure 13:**
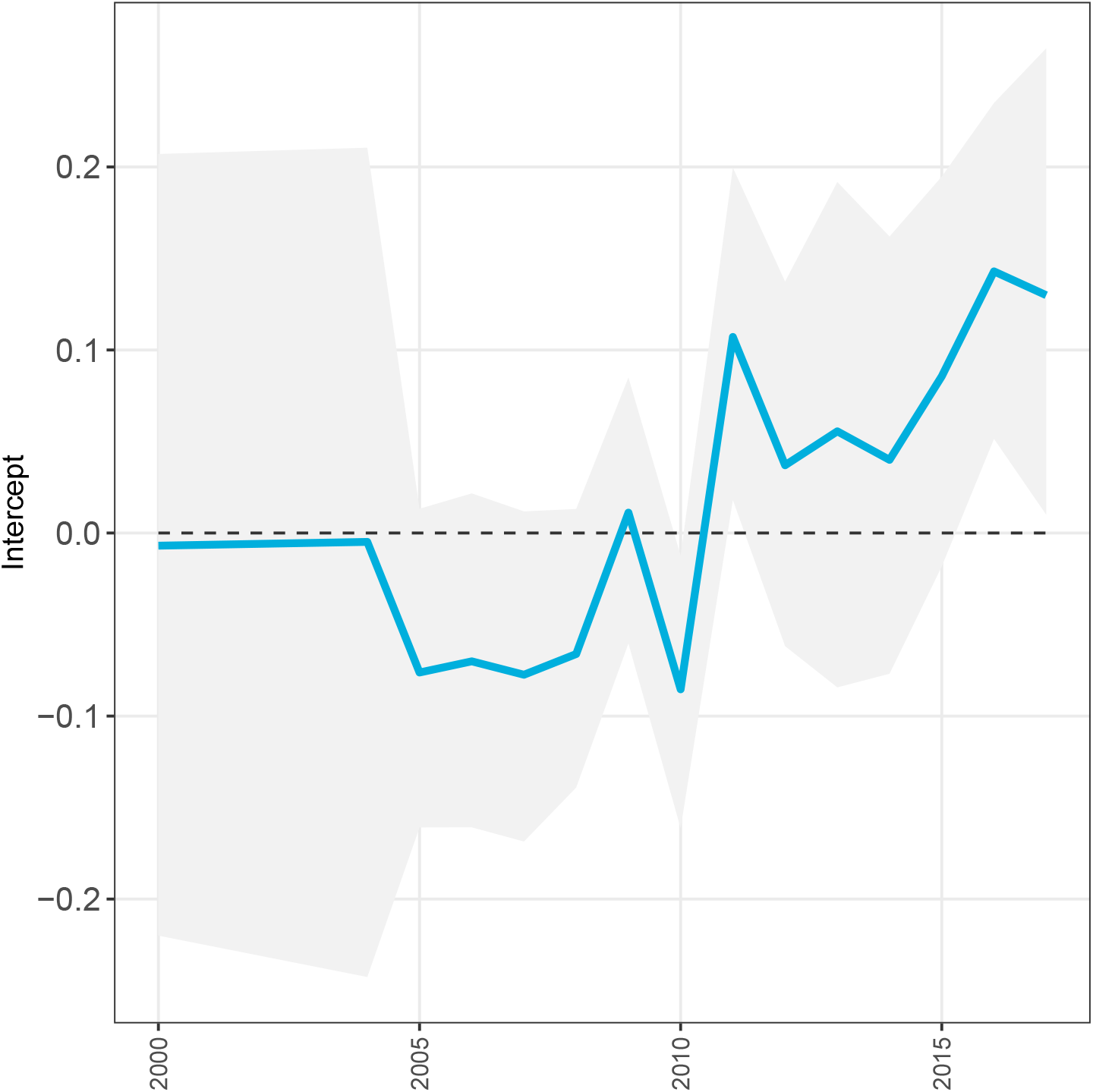
Random intercept values for each year in the detection model for wildlife plague risk.

**Extended Data Figure 14:**
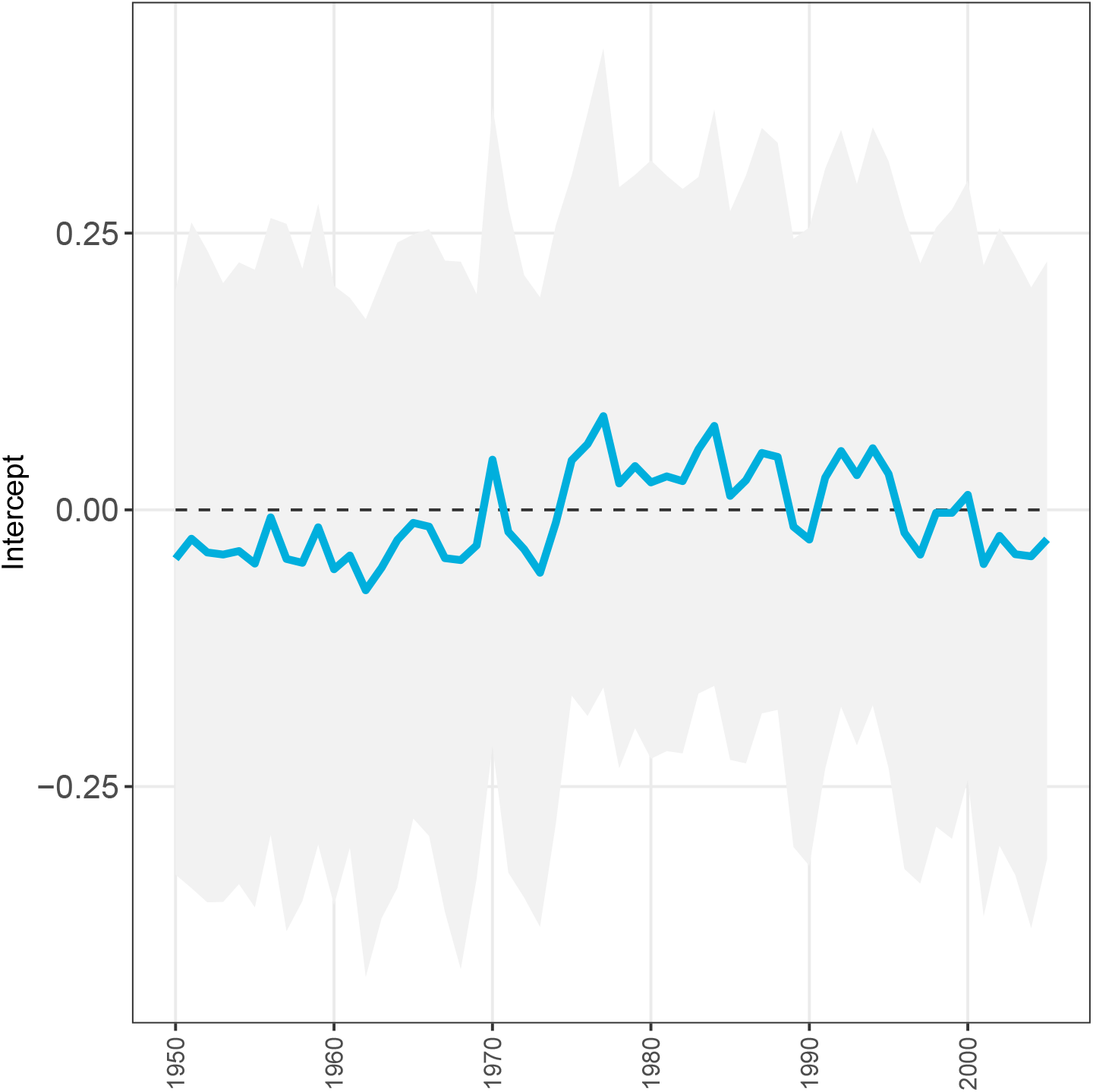
Random intercept values for each year in the detection model for human plague risk.

**Extended Data Figure 15:**
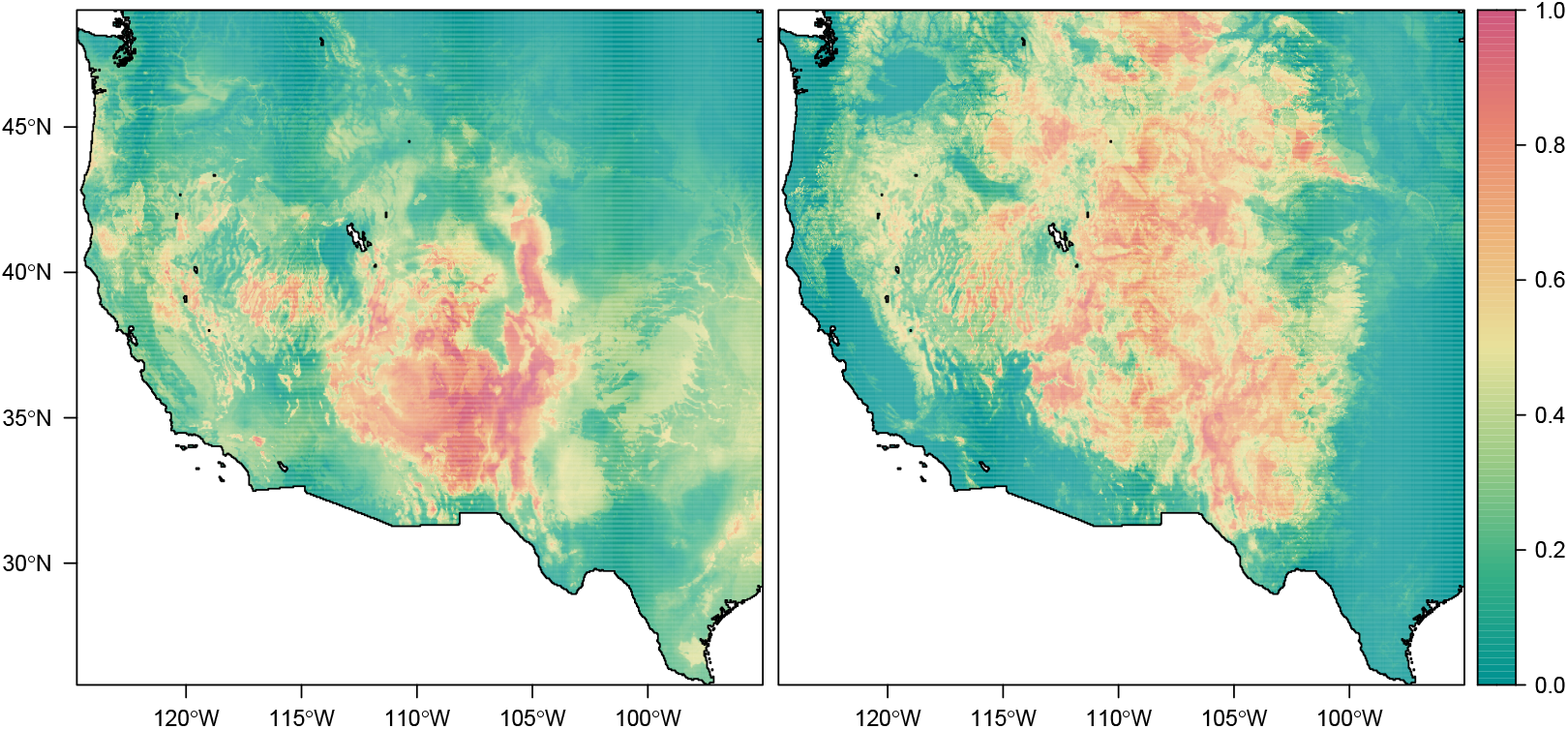
Mean environmental suitability across years in the detection model (left: human; right: wildlife). Random intercept models have nearly identical predictions to baseline models; the two are nearly perfectly correlated for humans (*r* = 0.992) and wildlife (*r* = 0.969).

**Extended Data Figure 16:**
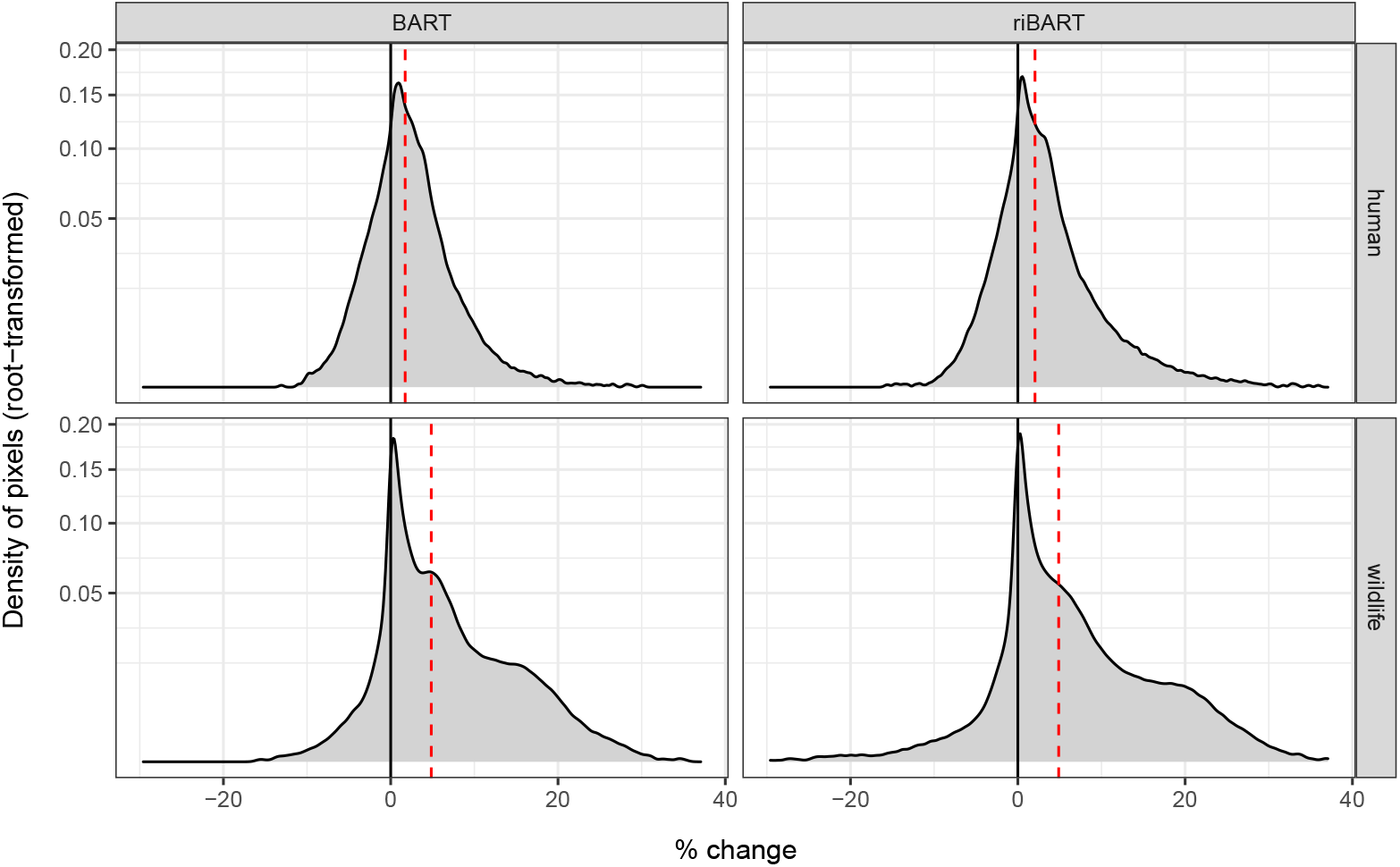
Estimated percent change in plague suitability, 1950 to present, across pixels and the four main models. Red lines show the mean value.

**Extended Data Figure 17:**
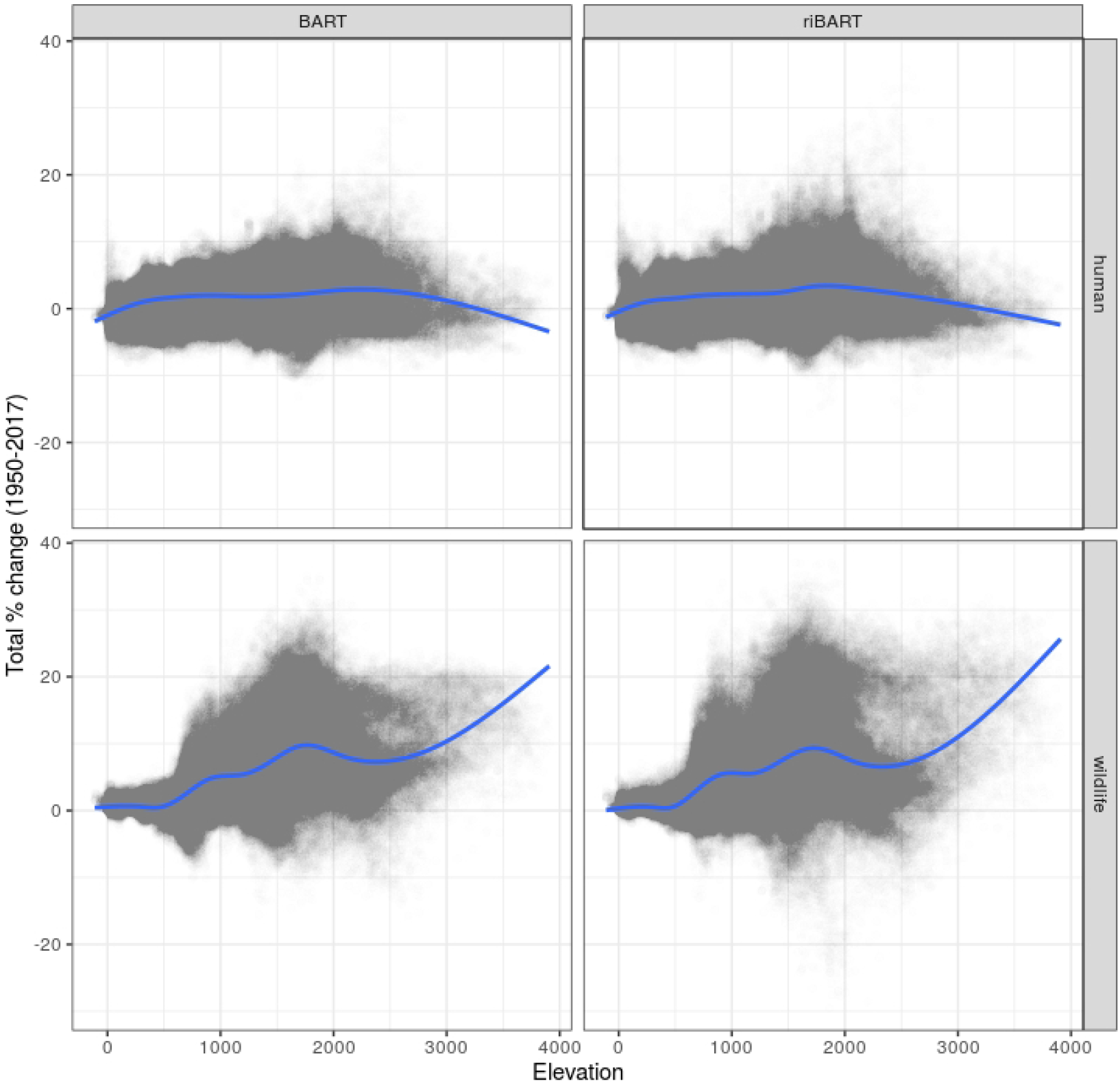
Estimated percent change in plague suitability, 1950 to present, versus elevation.

**Extended Data Figure 18:**
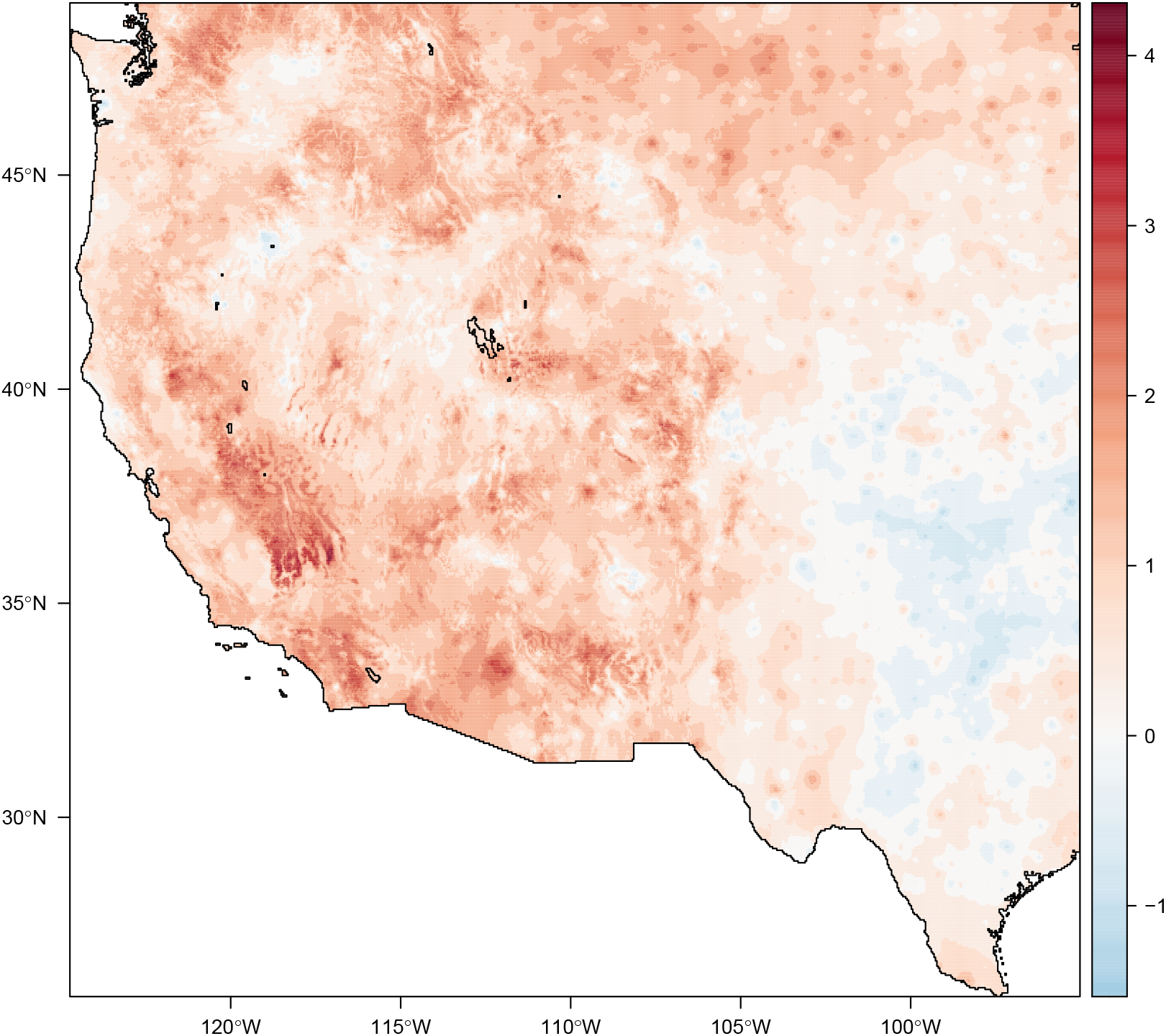
Total estimated change in mean temperature, 1950 to present; the region as a whole experienced an average warming of 0.84 °C.

**Extended Data Figure 19:**
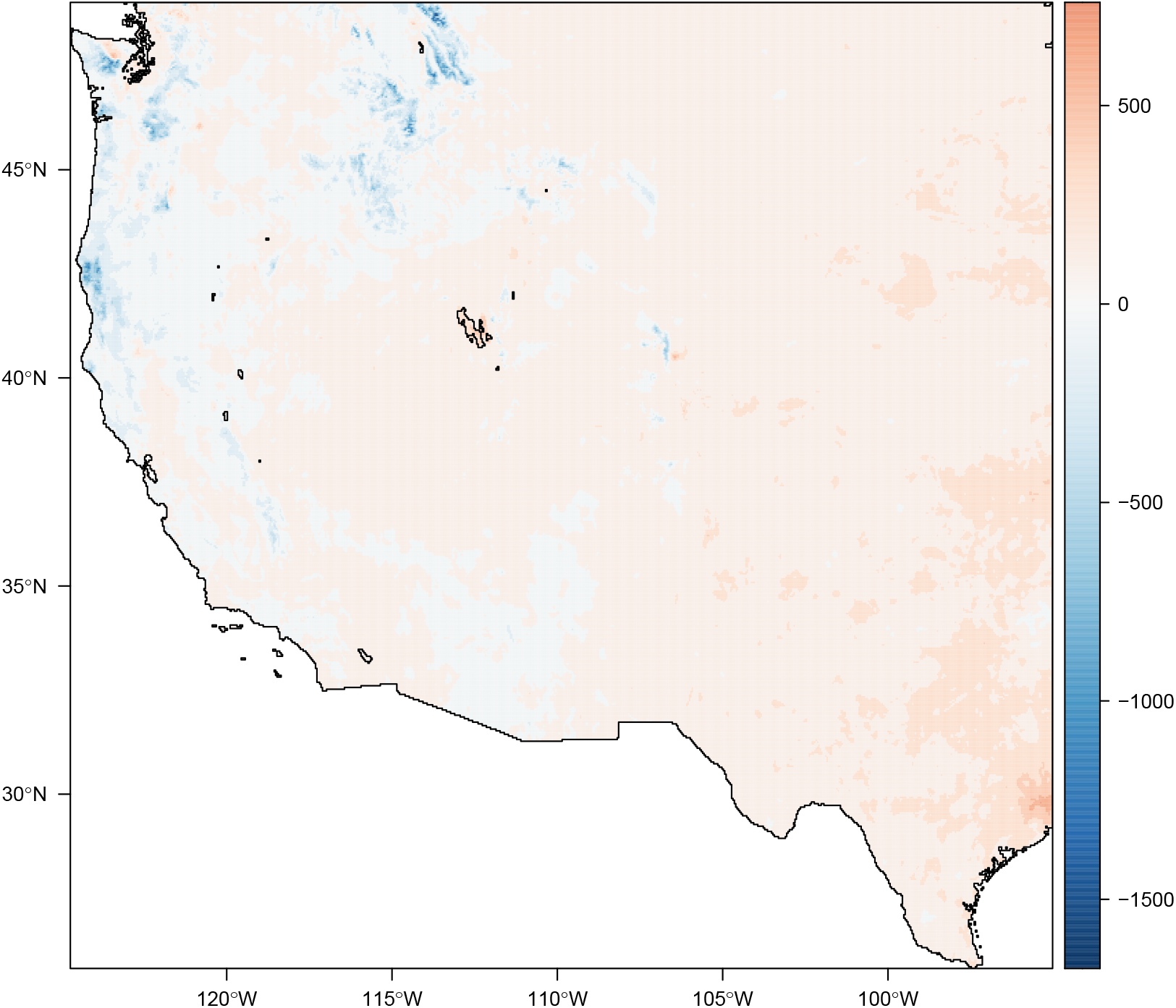
Total estimated change in precipitation, 1950 to present; the region as a whole experienced an average increase of 41.7 mm of annual precipitation.

**Extended Data Figure 20:**
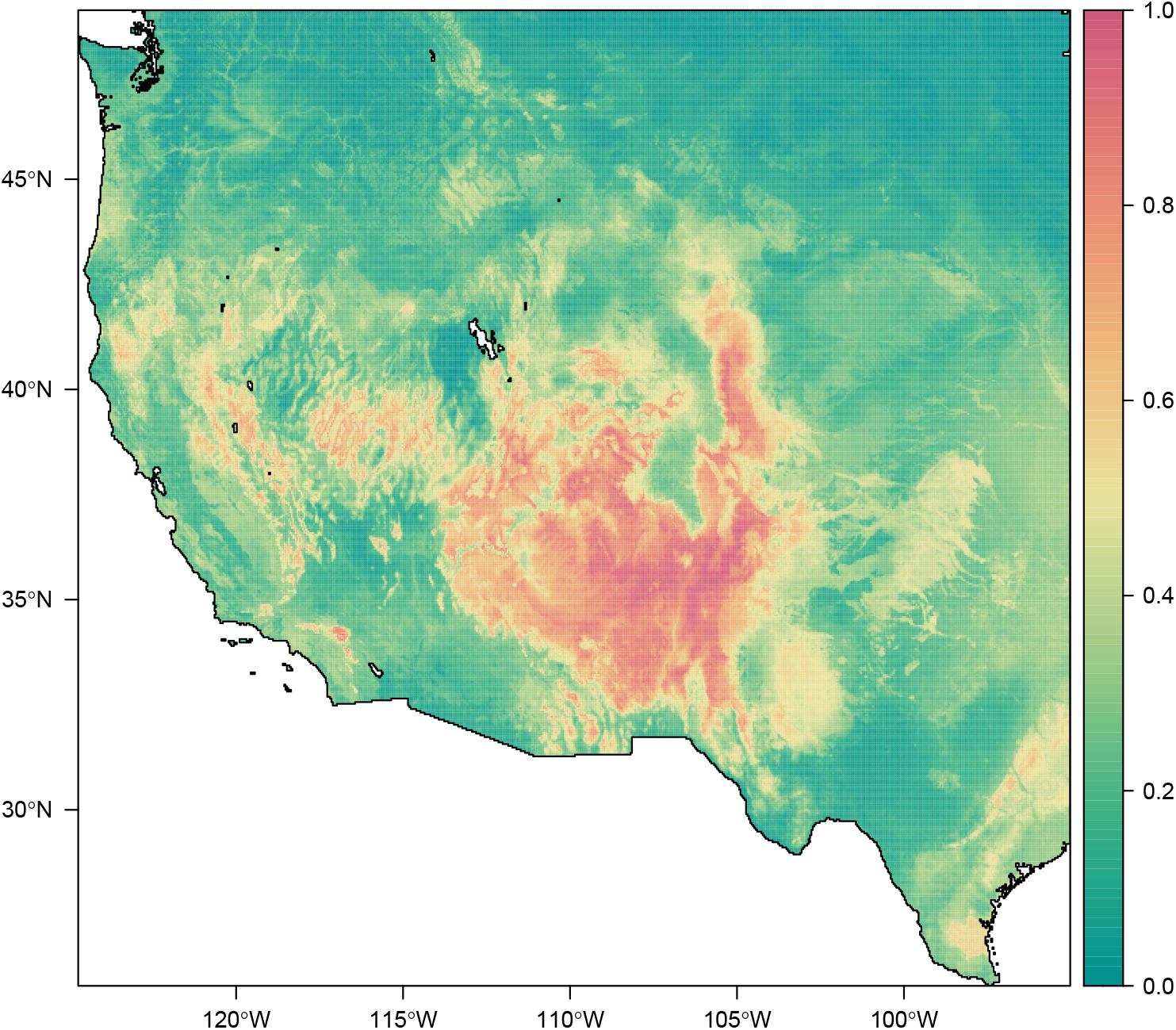
Mean suitability for plague across all years (1950-2017), using the alternate human model (no variable set reduction).

**Extended Data Figure 21:**
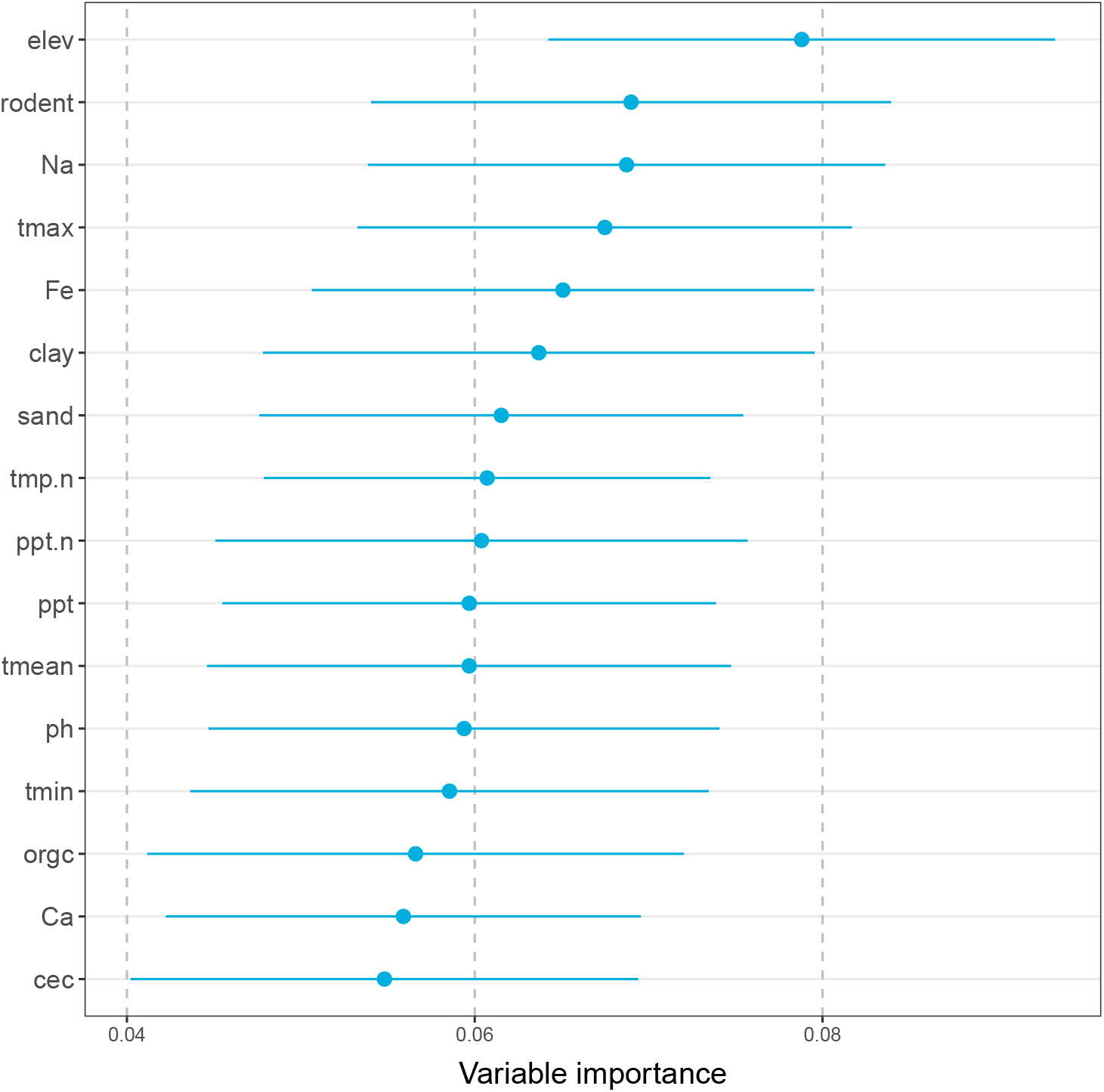
The variable importance diagnostic for the alternate human model using the full variables set (no variable set reduction).

**Extended Data Figure 22:**
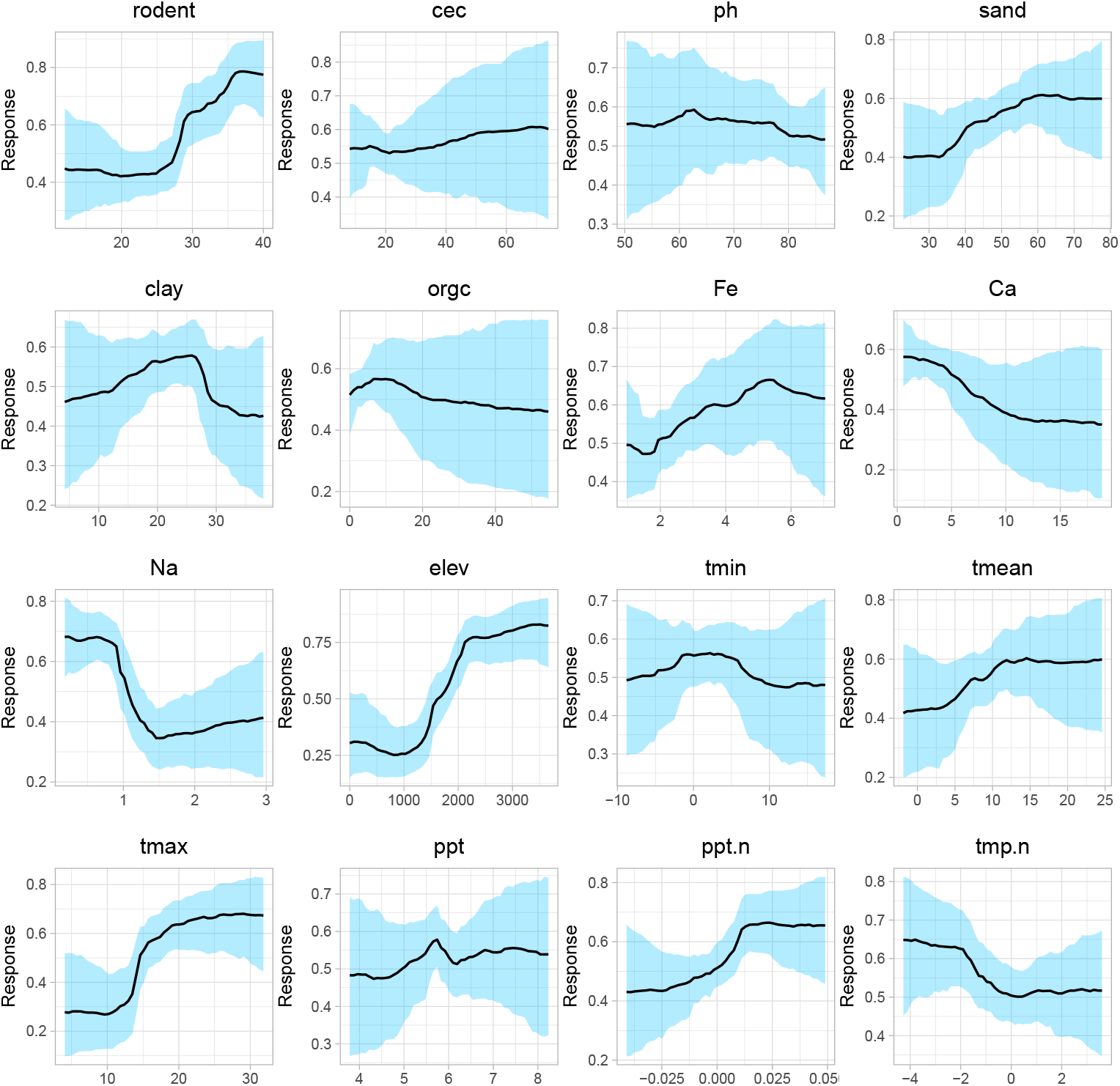
Partial dependence plots for the alternate human model using the full variables set (no variable set reduction).

**Extended Data Figure 23:**
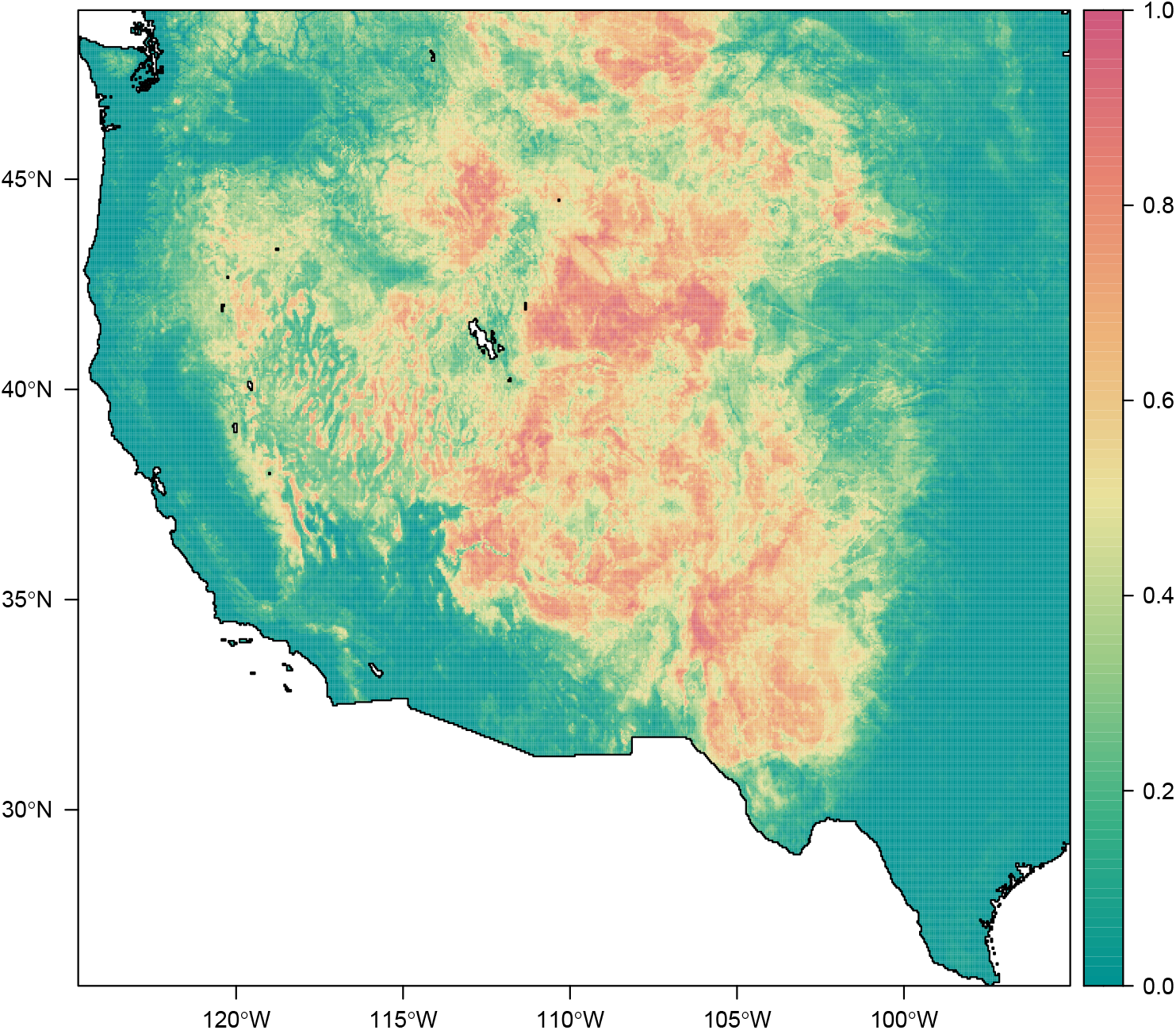
Mean suitability for plague across all years (1950-2017), using the first alternate wildlife model (no variable set reduction).

**Extended Data Figure 24:**
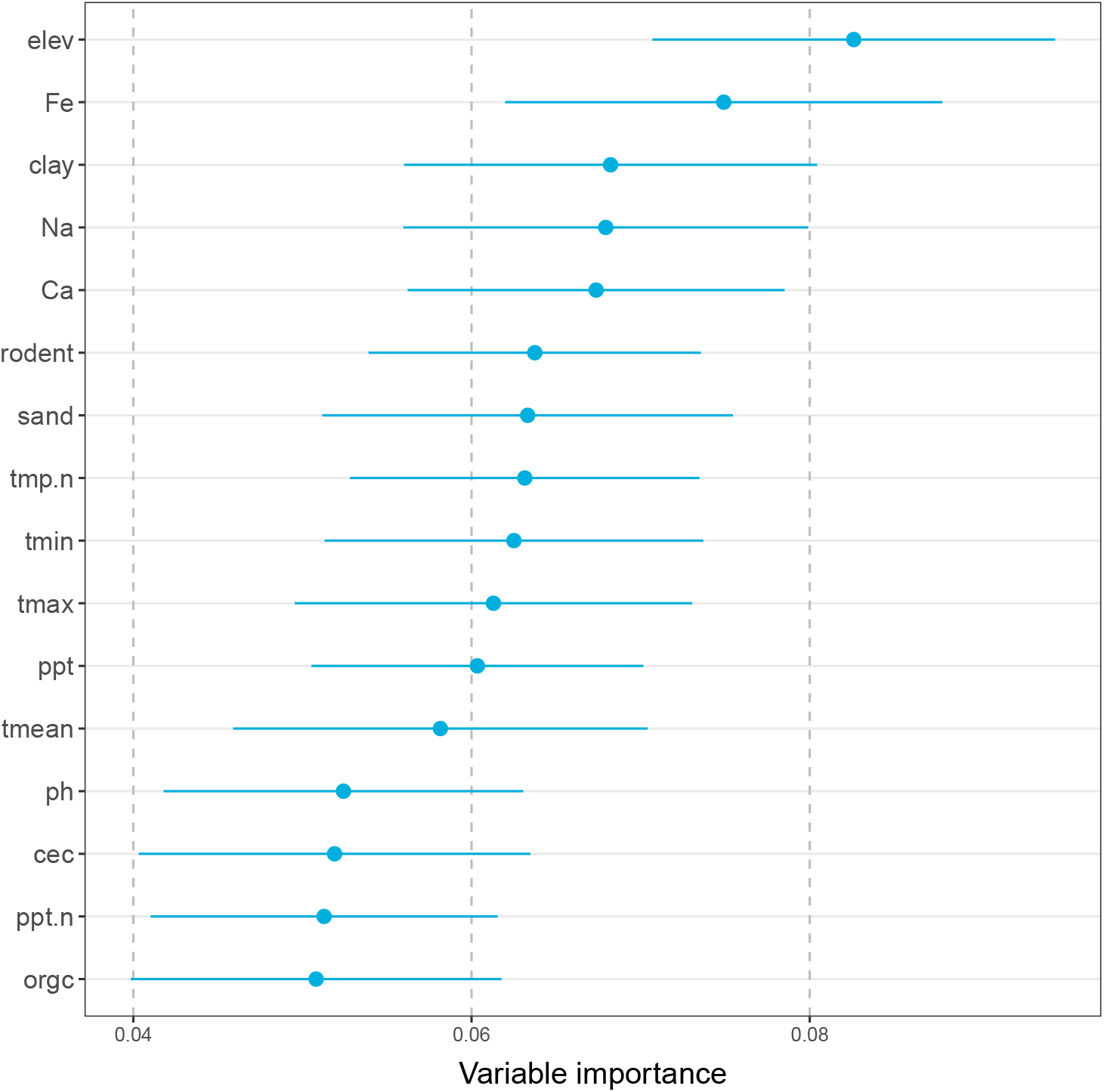
The variable importance diagnostic for the first alternate wildlife model using the full variables set (no variable set reduction).

**Extended Data Figure 25:**
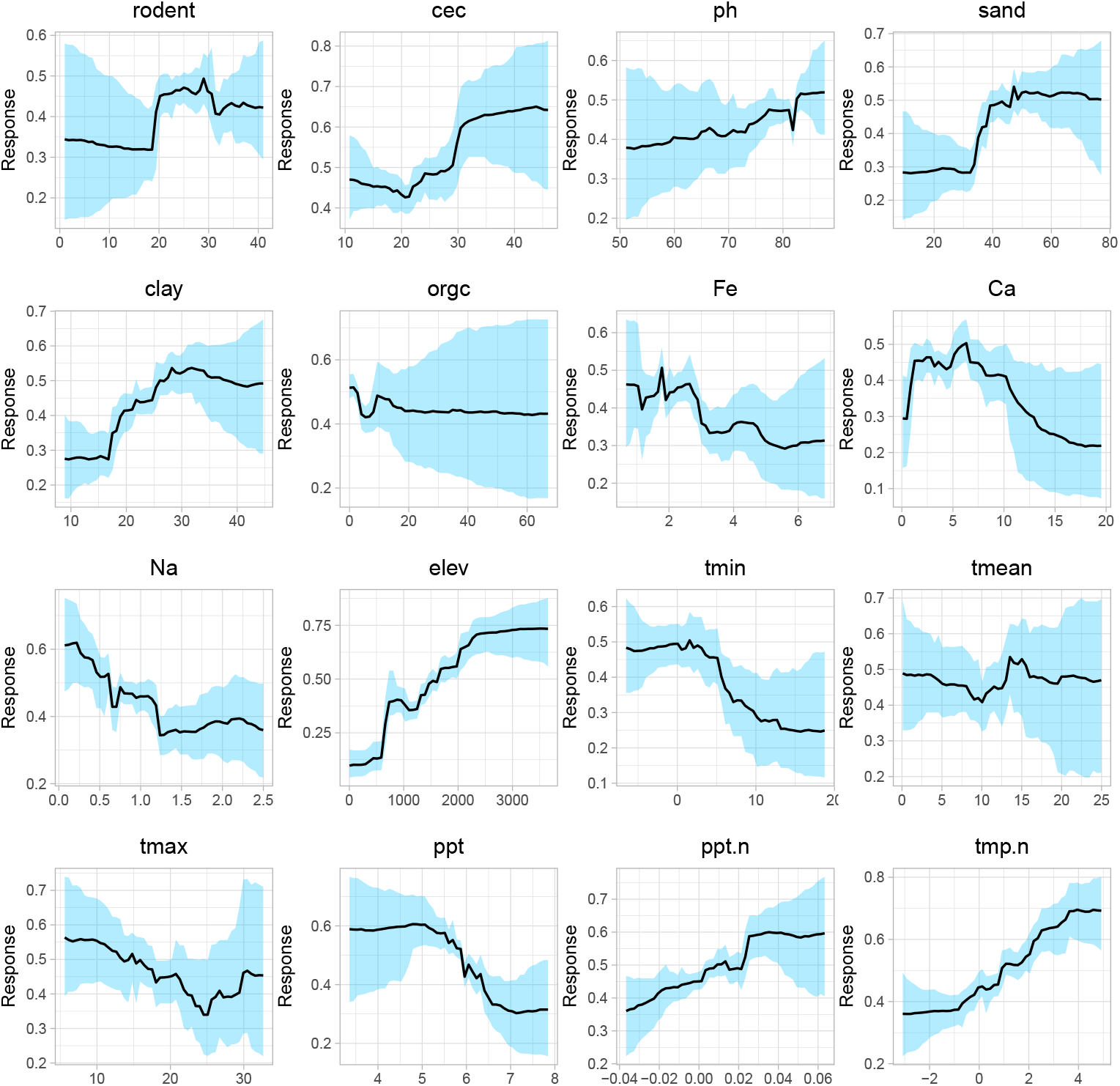
Partial dependence plots for the alternate wildlife model using the full variables set (no variable set reduction).

**Extended Data Figure 26:**
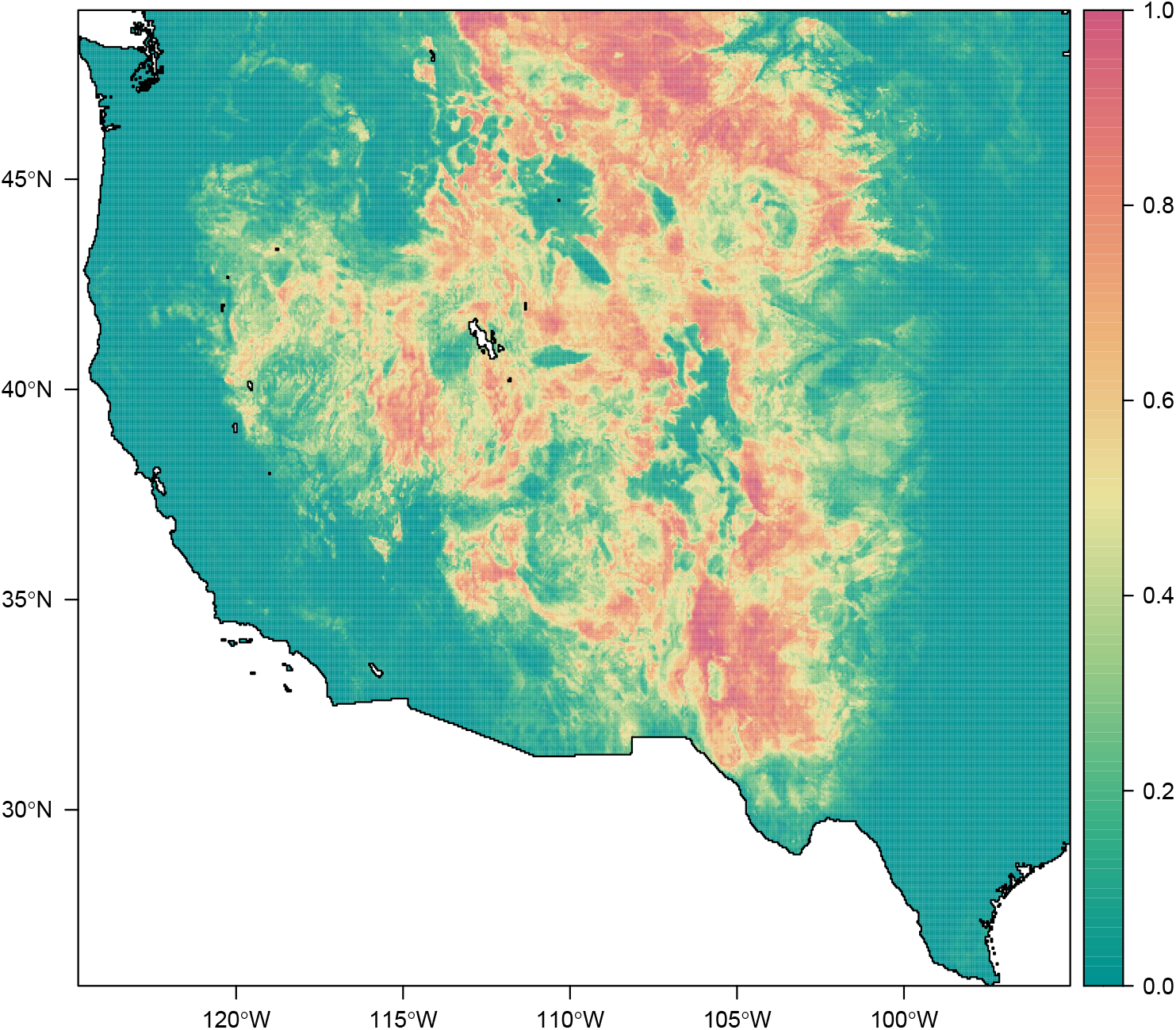
Mean suitability for plague across all years (1950-2017), using the second alternate wildlife model (pseudoabsences instead of true absences).

**Extended Data Figure 27:**
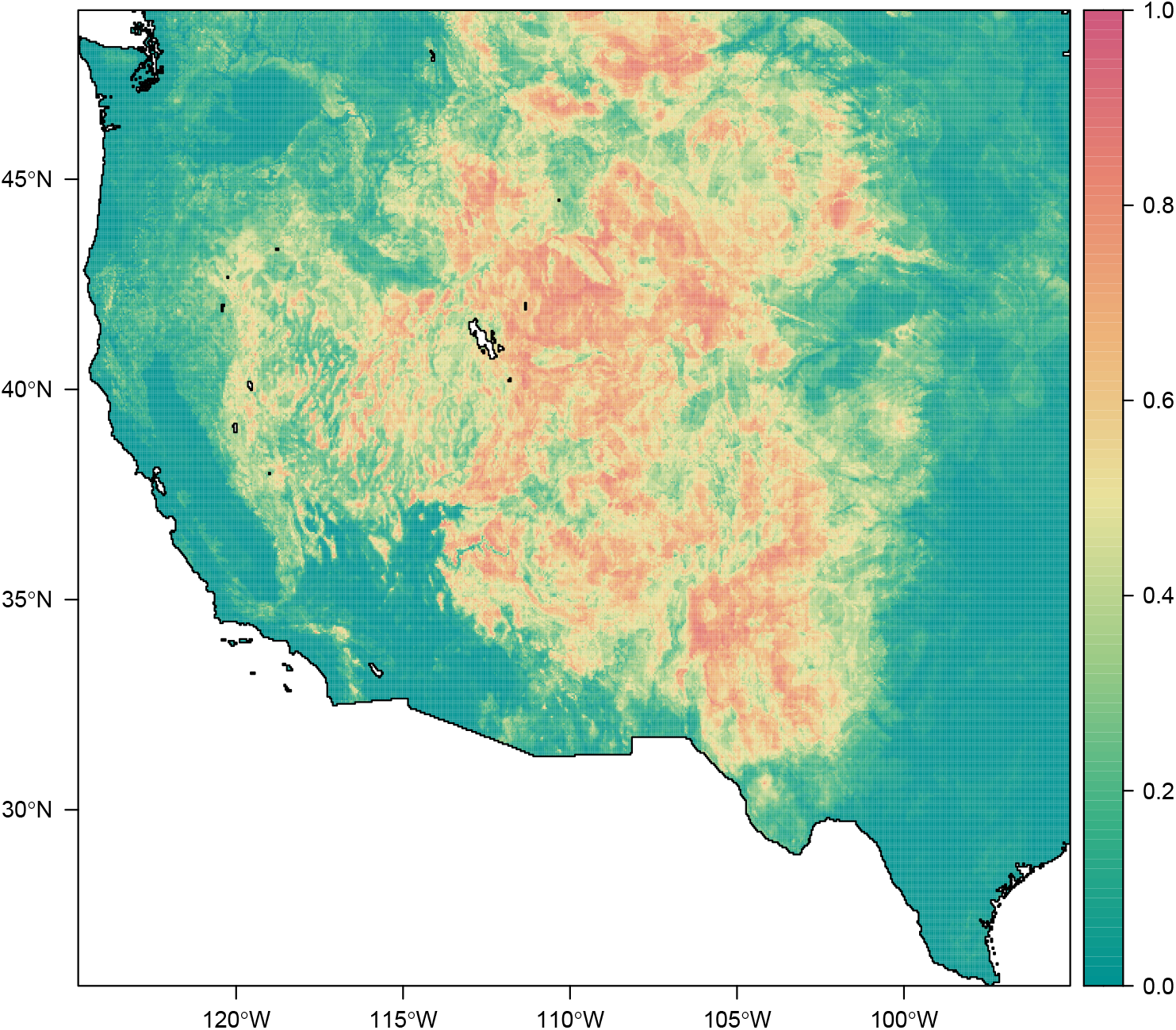
Mean suitability for plague across all years (1950-2017), using the second alternate wildlife model (only coyote data).

**Extended Data Table 1:** Variable name abbreviations used in this study, with full variable names and descriptions.

| **Variable** | **Full variable name** |
| --- | --- |
| rodent | Rodent species richness |
| cec | Soil cation exchange capacity |
| ph | Soil pH (acidity) |
| sand | Soil percent sand content by volume |
| clay | Soil percent clay content by volume |
| org | Soil organic carbon content |
| Fe | Soil iron macronutrient concentration |
| Ca | Soil calcium macronutrient concentration |
| Na | Soil sodium macronutrient concentration<br>(≈ proxy for salinity) |
| elev | Elevation above sea level |
| tmin | Average minimum annual temperature |
| tmean | Mean annual temperature |
| tmax | Average maximum annual temperature |
| ppt | Mean annual precipitation |
| ppt.n | Annual normalized precipitation anomaly<br>(relative to long-term average) |
| tmp.n | Annual normalized temperature anomaly<br>(relative to long-term average) |

**Extended Data Table 2:** Methodologies of previous studies applying ecological niche modeling to map plague (*Yersinia pestis*); this excludes studies focused on mapping individual reservoirs or fleas without any plague data. Abbreviations: PET/AET = potential and actual evapotranspiration; NDVI = normalized difference vegetation index; EVI = enhanced vegetation index; CTI = compound topographic index. ^†^ Note that this study also bootstrapped county level human cases in the U.S.

| Study | Extent | Years | Algorithm | Predictors |
| --- | --- | --- | --- | --- |
| 15 <sup>†</sup> | USA | 1965-2003? | GARP | Precipitation, temperature (min, mean, max), PET, AET, moisture surplus, moisture deficit, slope, aspect, elevation, CTI |
| 42 | Africa | 1970-2007 | GARP | BIO 1, 2, 5, 6, 12-14, slope, aspect, elevation, CTI, soil pH, soil moisture, soil carbon, PET, AET, humidity, growing degree days, NDVI |
| 54 | California, USA | 1984-2004 | MaxEnt | BIO 5, 7, 15-18 |
| 109 | “North America” | Unspecified | Maxent | BIO 1, 2, 5, 6, 12-14 |
| 59 | Western Usambara Mountains, Tanzania | 1986-2003 | GARP | EVI (mean, dry/rainy period means, standard error, seasonality, heterogeneity), slope, aspect, elevation, CTI |
| 63 | Northeast Brazil | 1966-2011 | GARP | BIO 1, 2, 5, 6, 12-14, NDVI, elevation |
| 110 | China | 1772-1964 | GARP | BIO 1, 2, 3, 7, 12, 14, 15 |
| 58 | Qinghai-Tibetan Plateau, China | 2004-2010 | MaxEnt | Elevation, land surface temperature (day & night), NDVI, slope, aspect, land cover |
| 57 | Western USA | 2000-2015 | MaxEnt | BIO 1, 11, 16, 17, distances to grassland, shrubland, cropland, sparse vegetation, bare soil, artificial surface, probability of deer mouse, altitude |

